# Extensive gene duplication in Arabidopsis revealed by pseudo-heterozygosity

**DOI:** 10.1101/2021.11.15.468652

**Authors:** Benjamin Jaegle, Rahul Pisupati, Luz Mayela Soto-Jiménez, Robin Burns, Fernando A. Rabanal, Magnus Nordborg

## Abstract

**Background:** It is apparent that genomes harbor massive amounts of structural variation, and that this variation has largely gone undetected for technical reasons. In addition to being inherently interesting, structural variation can cause artifacts when short-read sequencing data are mapped to a reference genome. In particular, spurious SNPs (that do not show Mendelian segregation) may result from mapping of reads to duplicated regions. Calling SNP using the raw reads of the 1001 Arabidopsis Genomes Project we identified 3.3 million heterozygous SNPs (44% of total). Given that *Arabidopsis thaliana* (*A. thaliana*) is highly selfing, we hypothesized that these SNPs reflected cryptic copy number variation, and investigated them further.

**Results:** The heterozygosity we observed consisted of particular SNPs being heterozygous across individuals in a manner that strongly suggests it reflects shared segregating duplications rather than random tracts of residual heterozygosity due to occasional outcrossing. Focusing on such pseudo-heterozygosity in annotated genes, we used GWAS to map the position of the duplicates, identifying 2500 putatively duplicated genes. The results were validated using *de novo* genome assemblies from six lines. Specific examples included an annotated gene and nearby transposon that, in fact, transpose together. Finally, we use existing bisulfite sequencing data to demonstrate that cryptic structural variation can produce highly inaccurate estimates of DNA methylation polymorphism.

**Conclusions:** Our study confirms that most heterozygous SNPs calls in *A. thaliana* are artifacts, and suggest that great caution is needed when analyzing SNP data from short-read sequencing. The finding that 10% of annotated genes exhibit copy-number variation, and the realization that neither gene- nor transposon-annotation necessarily tells us what is actually mobile in the genome suggest that future analyses based on independently assembled genomes will be very informative.

## Introduction

With the sequencing of genomes becoming routine, it is evident that structural variants (SVs) play a major role in genome variation (Alkan, Coe, and Eichler 2011). There are many kinds of SVs, e.g., indels, inversions, and transpositions. Of particular interest from a functional point of view is gene duplication, leading to copy number variation (CNV).

Before Next-Generation Sequencing (NGS) was available, genome-wide detection of CNVs was achieved using DNA-microarrays. These methods had severe weaknesses, leading to low resolution and problems detecting novel and rare mutations. (Carter 2007; Snijders et al. 2001). With the development of NGS, our ability to detect CNVs increased dramatically, using tools based on split reads, paired-end mapping sequencing coverage, or even *de novo* assembly (Shendure and Ji 2008; Zhao et al. 2013). In mammals, many examples of CNVs with a major phenotypic effect have been found (Gonzalez et al. 2005; Perry et al. 2007; Handsaker et al. 2011). One example is the duplication of MWS/MLS, associated with better trichromatic color vision (Miyahara et al. 1998).

While early investigation of CNV focused on mammals, several subsequent studies have looked at plant genomes. In *Brassica rapa*, gene CNV has been shown to be involved in morphological variation (Lin et al. 2014) and an analysis of the poplar “pan-genome” revealed at least 3000 genes affected by CNV (Pinosio et al. 2016). It has also been shown that variable regions in the rice genome are enriched in genes related to defense to biotic stress. (Yao et al. 2015). More recently, the first chromosome-level assemblies of seven accessions of *A. thaliana* based on long-read sequencing were released (Jiao and Schneeberger 2019), demonstrating that a large proportion of the genome is structurally variable. Similar studies have also been carried out in maize (C. Li et al. 2020; Hufford et al. 2021), tomato (Alonge et al. 2020), rice (Zhou et al. 2020) and soybean (Y. Liu et al. 2020). These approaches are likely to provide a more comprehensive picture than short-read sequencing, but are also far more expensive.

In 2016, the 1001 Genomes Consortium released short-read sequencing data and SNP calls for 1135 *A. thaliana* accessions (1001 Genomes Consortium 2016). Several groups have used these data to identify large numbers of structural variants using split reads (Göktay, Fulgione, and Hancock 2020; Zmienko et al. 2020; D.-X. Liu et al. 2021). Here we approach this from a different angle. Our starting point is the startling observation that, when calling SNPs in the 1001 Genomes data set, we identified 3.3 million (44% of total) putatively heterozygous SNPs. In a highly selfing organism, this is obviously highly implausible, and these SNPs were flagged as spurious: presumably products of cryptic CNV, which can generate “pseudo-SNPs” (Ranade et al. 2001; Hurles 2002) when sequencing reads from non-identical duplicates are (mis-)mapped to a reference genome that does not contain the duplication. Note that allelic SNP differences are expected to exist *ab initio* in the population, leading to instant pseudo-heterozygosity as soon as the duplicated copy recombines away from its template. In this paper we return to these putative pseudo-SNPs and show that they are indeed largely due to duplications, the position of which can be precisely mapped using GWAS. Our approach is broadly applicable, and we demonstrate that it can reveal interesting biology.

## Results

### Massive pseudo-heterozygosity in the 1001 Genomes data

Given that *A. thaliana* is highly selfing, a large fraction (44%) of heterozygous SNPs is inherently implausible. Two other lines of evidence support the conclusion that they are spurious. First, genuine residual heterozygosity would appear as large genomic tracts of heterozygosity in individuals with recent outcrossing in their ancestry. Being simply a random product of recombination and Mendelian segregation, there is no reason two individuals would share tracts unless they are very closely related. The observed pattern is completely the opposite. While a small number of individuals do show signs of recent outcrossing, this is quite rare (as expected given the low rate of outcrossing in this species, and the fact that the sequenced individuals were selected to be completely inbred). Instead we find that the same SNP are often heterozygous in multiple individuals. Although the population level of heterozygosity at a given SNP is typically low **(Supplemental Figure 1)**, over a million heterozygous SNPs are shared by at least 5 accessions, and a closer look at the pattern of putative heterozygosity usually reveals short tracts of shared heterozygosity that would be vanishingly unlikely under residual heterozygosity, but would be expected if tracts represent shared duplications, and heterozygosity is, in fact, pseudo-heterozygosity due to mis-mapped reads (**Figure 1**). Analysis of the distribution of the lengths and number of putatively heterozygous tracts across accessions shows that the vast majority of accessions have a large number of very short tracts (roughly 1 kb) of heterozygosity (**Supplemental Figure 2**). Longer tracts are rare and not shared between accessions.

**Figure 1:**
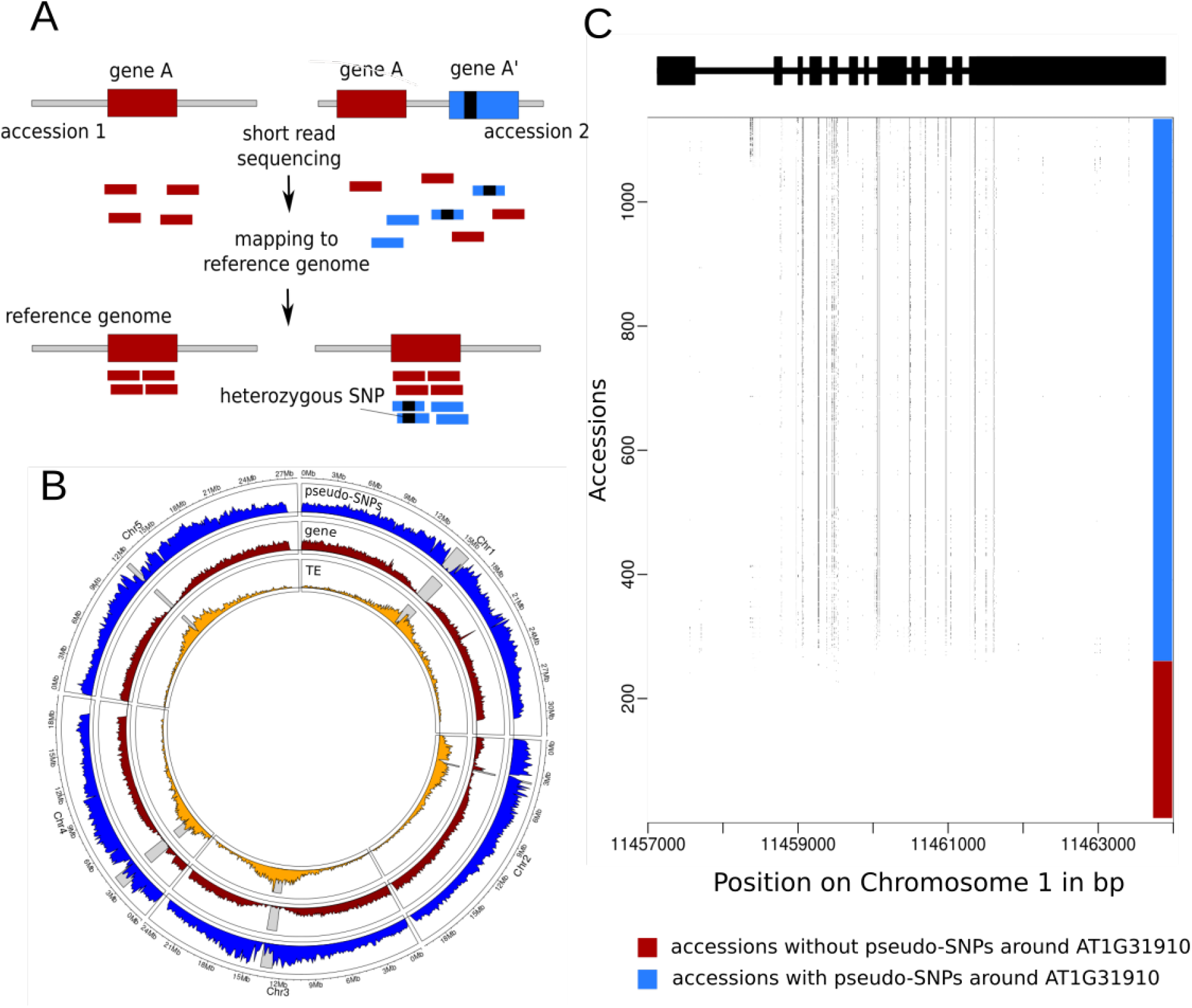
Pseudo-heterozygosity in the 1001 Genomes dataset. (**A**) Cartoon illustrating how a duplication can generate pseudo-SNPs when mapping to a reference genome that does not contain the duplication. (**B**) Genomic density of transposons, genes, and shared heterozygous SNPs. Gray bars represent the position of the centromere for each chromosome. (**C**) The pattern of putative heterozygosity around AT1G31910 for the 1057 accessions. Dots in the plot represent putative heterozygosity.

Furthermore, the density of shared heterozygous SNPs is considerably higher around the centromeres (**Figure 1**), which is again not expected under random residual heterozygosity, but is rather reminiscent of the pattern observed for transposons, where it is interpreted as the result of selection removing insertions from euchromatic regions, leading to a build-up of common (shared) transposon insertions near centromere (Quadrana et al. 2016). As we shall see below, it is likely that transposons play an important role in generating cryptic duplications leading to pseudo-heterozygosity (although we emphasize again that the heterozygous SNPs were called taking known repetitive sequences into account).

Despite the evidence for selection against these putative duplications, we found 2570 genes containing 26647 pseudo-SNPs segregating at 5% or more in the population (**Supplemental Figure 3**). Gene-ontology analysis of these genes reveals an enrichment for biological processes involved in response to UV-B, bacteria or fungi (**Supplemental Figure 4**). In the following sections, we investigate these putatively duplicated genes further.

### Mapping common duplications using genome-wide association

If heterozygosity is caused by the presence of cryptic duplications in non-reference genomes, it should be possible to map the latter using GWAS with heterozygosity as a “phenotype” (Imprialou et al 2017). We did this for each of the 26647 SNPs exhibiting shared heterozygosity within genes (**Supplemental Figure 3**).

Of the 2570 genes that showed evidence of duplication, 2511 contained at least one major association (using significance threshold of *p* < 10^−20^; see Methods). For 708 genes, the association was more than 50 kb away from the pseudo-SNP used to define the phenotype, and for 175 it was within 50 kb. We will refer to these as *trans-* and *cis-*associations, respectively. The majority of genes, 1628, had both *cis-* and *trans-*associations (**Figure 2**).

**Figure 2:**
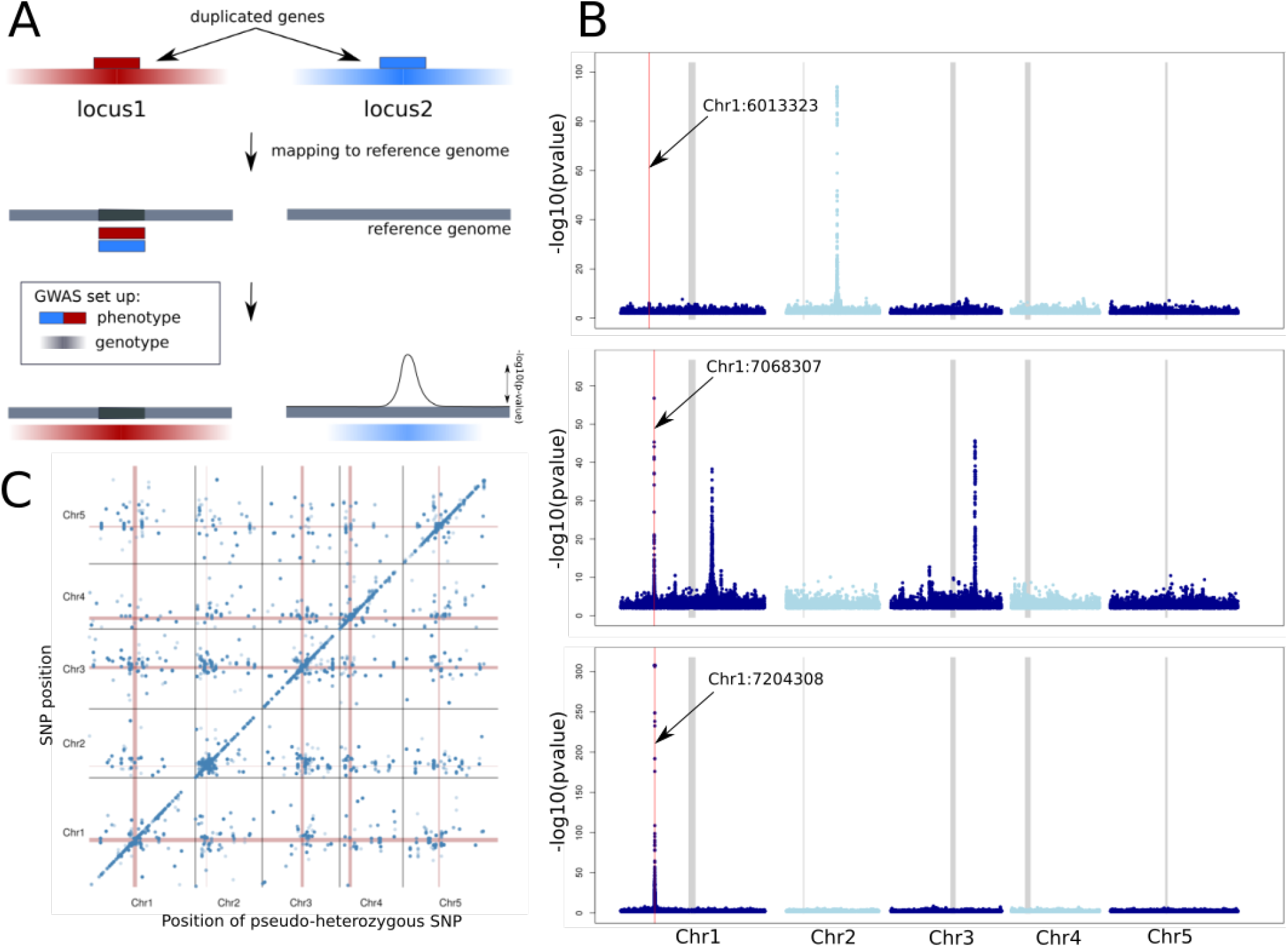
GWAS of putative duplications **(A)** Schematic representation of the principle of how GWAS can be used to detect the position of the duplicated genes based on linkage disequilibrium (LD). As phenotype, heterozygosity at the position of interest is coded as 1 (present) or 0 (absent). As genotype, the SNPs matrix of the 1001 genome dataset was used (with heterozygous SNPs filtered out). Color gradients represent the strength of LD around the two loci. In this example the reference genome does not contain locus2. (**B**) GWAS results for three different genes with evidence of duplication, for illustration. The red lines indicate the position of the pseudo-SNP used for each gene/GWAS and the thick grey lines indicate the centromeres. The top plot shows a *trans*-association, the bottom a *cis*-association, and the middle shows a case with both (*cis* plus two *trans*). (**C**) Summary of all 26647 GWAS results.

To validate these results, we assembled 6 non-reference genomes *de novo* using long-read PacBio sequencing. The GWAS results provide predicted locations of the duplications (the putative causes of pseudo-heterozygosity). We identified the homologous region of each non-reference genome, then used BLAST to search for evidence of duplication. For 84% of the 403 genes predicted to have a duplication present in at least one of the six non-reference genomes, evidence of a duplication was found; for 60%, the occurrence perfectly matched the pattern of heterozygosity across the six genomes. For the remaining 16%, no evidence of a duplication was found, which could be due to the stringent criteria we used to search for evidence of duplication (see Methods). The distribution of fragment sizes detected suggests that we capture a mixture of duplicated gene fragments and full genes (**Supplemental Figure 5**).

### Rare duplications

The GWAS approach has no power to detect rare duplications, which is why we restricted the analysis above to pseudo-heterozygous SNPs seen in five or more individuals. Yet most are rarer: 40% are seen only in a single individual, and 16% are seen in two. As it turns out, many of these appear to be associated with more common duplications. Restricting ourselves to genes only, 11.4% of the singleton pseudo-heterozygous SNPs are found in the 2570 genes already identified using common duplications, a significant excess (*p* = 2.5e-109). For doubletons, the percentage is 11.1% (*p* = 1.9e-139). Whether they are caused by the same duplications, or reflect additional ones present at lower frequency is difficult to say. To confirm duplications more directly, we took the reads generating the singleton and doubleton pseudo-heterozygotes, and compared the result of mapping them to the reference genome, and to the appropriate genome (derived from the same inbred line). One predicted consequence of the reads mapping at different locations is that mapping coverage around the pseudo-SNPs will be decreased when mapping to the newly assembled PacBio genomes rather than the reference genome. As expected, a high proportion of the SNPs tested have lower coverage when mapping to the PacBio genomes **(Supplemental Figure 6-7)**. In addition to a decrease in coverage, we were also able to detect reads mapping to multiple locations in the right genomes, as well as the corresponding disappearance of the pseudo-SNPs. For example, 41.5% of the doubletons tag regions that map to more regions in the PacBio genomes than in the reference genome (**Supplemental Figure 6-8**).

### Local duplications

If duplications arise via tandem duplications, they will not give rise to pseudo-SNPs until the copies have diverged via mutations. This is in contrast to unlinked copies, which will lead to pseudo-SNPs due to existing allelic variation as soon as recombination has separated copy from original. We should thus expect the approach taken here to be biased against detecting local duplications. Nonetheless, GWAS revealed 175 genes with evidence only for a *cis* duplication. 28 of these were predicted to be present in at least one of the six new genomes, and 14 could be confirmed to have local variation of copy number relative to the reference. (**Figure 3A**).

**Figure 3:**
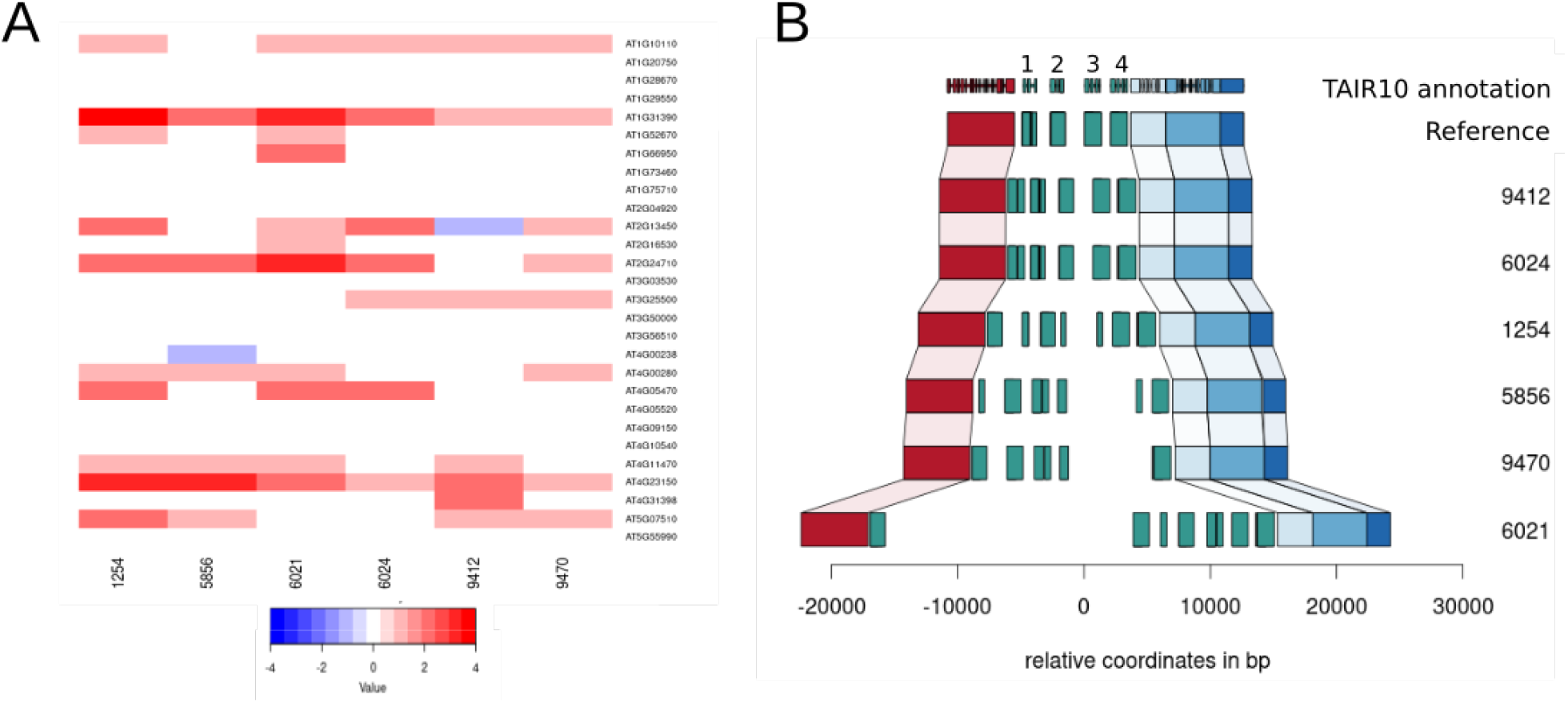
Confirmation of tandem duplications. (**A**) The distribution of estimated copy number (based on sequencing coverage) across 6 PacBio genomes for 28 genes predicted to be involved in tandem duplications based on the analyses of this paper. (**B**) The duplication pattern observed in these genomes for the gene AT1G31390, as an example The reference genome contains four copies, shown as numbered green boxes. Other colored boxes denote other genes.

The local structure of the duplications can be complex. An example is provided by the gene AT1G31390, annotated as a member of MATH/TRAF-domain genes, and which appears to be present in 4 tandem copies in the reference genome, but which is highly variable between accessions, with one of our accessions carrying at least 6 copies (**Figure 3B)**. However, there are no copies elsewhere in any of the new genomes for this gene (**Supplemental Figure 9)**.

### Transposon-driven duplications

Transposons are thought to play a major role in gene duplications, capturing and moving genes or gene fragments around the genome (Woodhouse, Pedersen, and Freeling 2010; Lisch 2013). While confirming the *trans* duplications in the PacBio genomes, we found a beautiful example of this process. The gene AT1G20400 (annotated, based on sequence similarity, to encode a myosin heavy chain-like protein) was predicted to have multiple *trans*-duplications. The 944 bp coding region contains 125 putatively heterozygous SNPs with striking haplotype structure characteristic of structural variation **(Figure 4C)**. We were able to identify the duplication predicted by GWAS in the six new genomes **(Figure 4)**. Four of the newly assembled genomes have only one copy of the gene, just like the reference genome, but one has 3 copies and one has 4 copies. However, none of the 6 new genomes has a copy in the same place as in the reference genome (**Supplemental Figure 10**).

**Figure 4:**
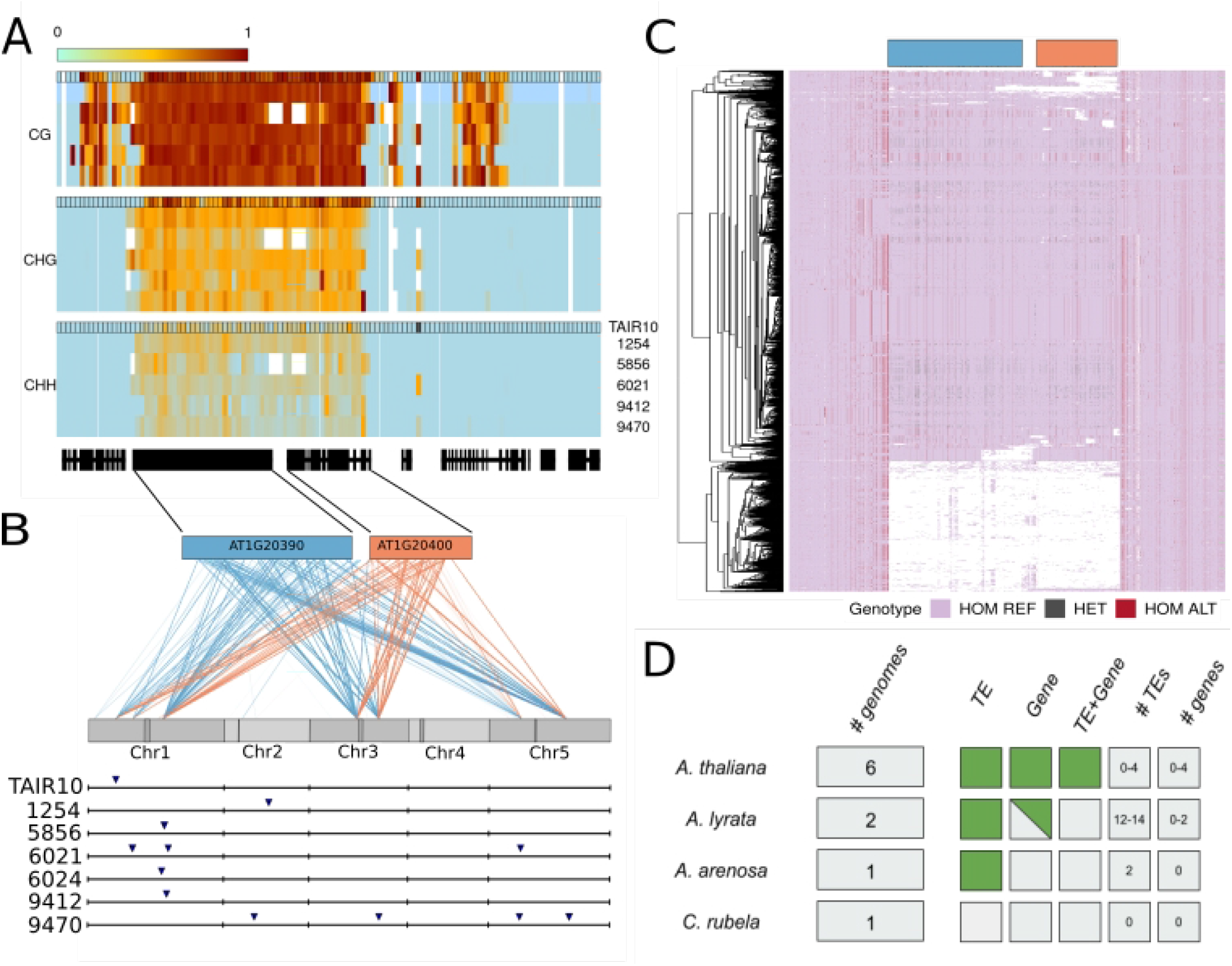
A Gypsy element (AT1G20390) and a gene transpose (AT1G20400) together. (**A**) Methylation levels on regions containing AT1G20390 and AT1G20400 for 6 accessions, calculated in 200 bp windows after mapping reads to the TAIR10 reference genome (annotation outline in black). (**B**) GWAS results for the putatively heterozygous SNPs in AT1G20390 and AT1G20400. Each line represents the link between the position of the pseudo-SNP and a GWAS hit position in the genome. The lower part shows the presence of the new transposable element in the 6 PacBio genomes as well as in the reference genome. (**C**) SNP haplotypes around the AT1G20400 region in the 1001 genomes data. White represents a lack of coverage. (**D**) Presence of the gene and the transposon in related species.

In the reference genome, AT1G20400 is closely linked to AT1G20390, which is annotated as a Gypsy element. This element also contains many pseudo-SNPs, and GWAS revealed duplication sites overlapping those for AT1G20400 (**Figure 4B**). This suggested that the putative gene and putative Gypsy element transpose together, i.e. that both are misannotated, and that the whole construct is effectively a large transposable element. Further analysis of the PacBio genomes confirmed that AT1G20400 and AT1G20390 were always found together, and we were also able to find conserved Long Terminal Repeat sequences flanking the whole construct, as would be expected for a retrotransposon (**Supplemental Figure 11-12**). We did not find any evidence for expression of AT1G20400 in RNAseq from seedlings in any of the accessions. Available bisulfite sequencing data (Kawakatsu et al. 2016) showed that the whole region is heavily methylated, as expected for a transposon **(Figure 4)**. We tried mapping the bisulfite reads to the appropriate genome for the respective accesions, but the coverage was too low and noisy to observe a difference in methylation between the multiple insertions (**Supplemental Figure 13)**.

Having located precise insertions in the six new genomes, we attempted to find them using short-read data in the 1001 Genomes dataset. Except for one insertion that was shared by 60% of accessions, the rest were found in less than 20%, suggesting that this new element has no fixed insertions in the genome — including the insertion found in the TAIR10 reference genome, which was only found in 17.4 % of the accessions (**Supplemental Figure 14**). We also looked for the element in the genomes of *A. lyrata* (two different genomes), *A. suecica* (a tetraploid containing an A. thaliana and an A. arenosa subgenome; see Burns et al. 2021), and *Capsella rubella (Slotte et al. 2013)*. The gene and the Gypsy element were only found together in *A. thaliana* (including the *A. thaliana* sub-genome of the allopolyploid *A. suecica*). The Gypsy element alone is present in the other *Arabidopis* species, and the gene alone is present in *A. lyrata*, but only in one of two genomes. In *Capsella rubella*, neither the transposon nor the gene could be detected **(Supplemental Figure 15)**. Thus the transposon and gene appears to be specific to the genus *Arabidopsis*, while their co-transposition is specific to *A. thaliana*, suggesting that the new transposable element evolved since divergence of *A. thaliana* from the other member of the genus.

### Spurious methylation polymorphism

Just like cryptic duplications can lead to spurious genetic polymorphisms, they can lead to spurious cytosine methylation polymorphisms. Indeed, given the well-established connection between gene duplication and gene silencing (e.g., Melquist, Luff, and Bender 1999), they may be more likely to do so. To investigate this, we re-examined the methylation status of genes previously reported by the 1001 Genomes Project (Kawakatsu et al. 2016) as having complex patterns of methylation involving both CG and CHG methylation. In our six sequenced accessions, we found 19530 genes that had been reported as having CG methylation (in at least one accession) and 2556 genes that had been reported as having CHG methylation (in at least one accession). 2473 genes were part of both sets. Out of these, 619, or 24%, had been detected as duplicated in the analyses presented above (a massive enrichment compared to the genome-wide fraction of roughly 10%). To understand these patterns better, we mapped the original bisulfite data to the appropriate genome as well as to the reference genome. In any given accession, roughly 7% of the 2473 genes could not be compared because the homologous copy could not be found (this is presumably mostly because they contain structural variants that prevent them being located by BLAST; see Supplementary Table 1), and roughly 30% exhibited copy number variation (Table 1). The remaining genes had a single match, almost always in the same location as in the reference genome. These categories are shared across accessions: 1294 of the 2367 genes appeared to be single-copy in all six new genomes, for example (Table 1; Additional files 1-8).

**Table 1.** Number of copies of the 2367 genes identified in each new genome (and Araport11, as control).

| Target | Number of copies identified |  |  |
| --- | --- | --- | --- |
|  | 0 | 1 | >1 |
| 1254 | 138 | 1563 | 772 |
| 5856 | 174 | 1566 | 733 |
| 6021 | 131 | 1577 | 765 |
| 6024 | 152 | 1554 | 767 |
| 9412 | 147 | 1567 | 759 |
| 9470 | 142 | 1589 | 742 |
| *Intersection* | *37* | *1294* | *610* |
| Araport11 | 0 | 1721 | 752 |

Turning to the methylation patterns, the effect of cryptic copy number variation was obvious (Table 2). For the genes with a single match in both the reference and accession genome, methylation status calls based on mapping bisulfite sequencing reads to either genome were largely concordant (roughly 2.5% disagreement), whereas for genes with copy number variation, roughly one third of calls were wrong.

**Table 2.** Fraction of differentially methylated genes when comparing bisulfite reads mapped to reference TAIR genome and to its respective PacBio genome, separated by gene copy number.

| Target | Number of copies identified |  |  |  |
| --- | --- | --- | --- | --- |
|  | 1 |  | >1 |  |
|  | CG (%) | CHG (%) | CG (%) | CHG (%) |
| 1254 | 3.0 | 4.4 | 33.3 | 21.6 |
| 5856 | 1.2 | 3.7 | 27.8 | 42.9 |
| 6021 | 2.4 | 3.2 | 39.3 | 24.2 |
| 6024 | 3.0 | 4.2 | 41.2 | 29.5 |
| 9412 | 2.0 | 2.5 | 37.0 | 27.1 |
| 9470 | 2.1 | 4.7 | 36.0 | 26.2 |

As an illustration for why this occurs, consider the methylation status of AT1G30140 (**Figure 5**). When mapped to the reference genome, 5 out of 6 accessions were found to be both CG and CHG methylated, with accession 6021 having no methylation. When mapped to the appropriate genome, we see that this pattern can be quite misleading. In accession 1254, for example, we found three apparent copies of the gene, only two of which are methylated, neither of which is the copy corresponding to the copy present in the reference genome. In accession 5856, the copy corresponding to the reference genome cannot be identified, but a copy on a different chromosome is identified, and it is methylated. In both cases, mapping to the reference genome leads to incorrect methylation status for AT1G30140.

**Figure 5:**
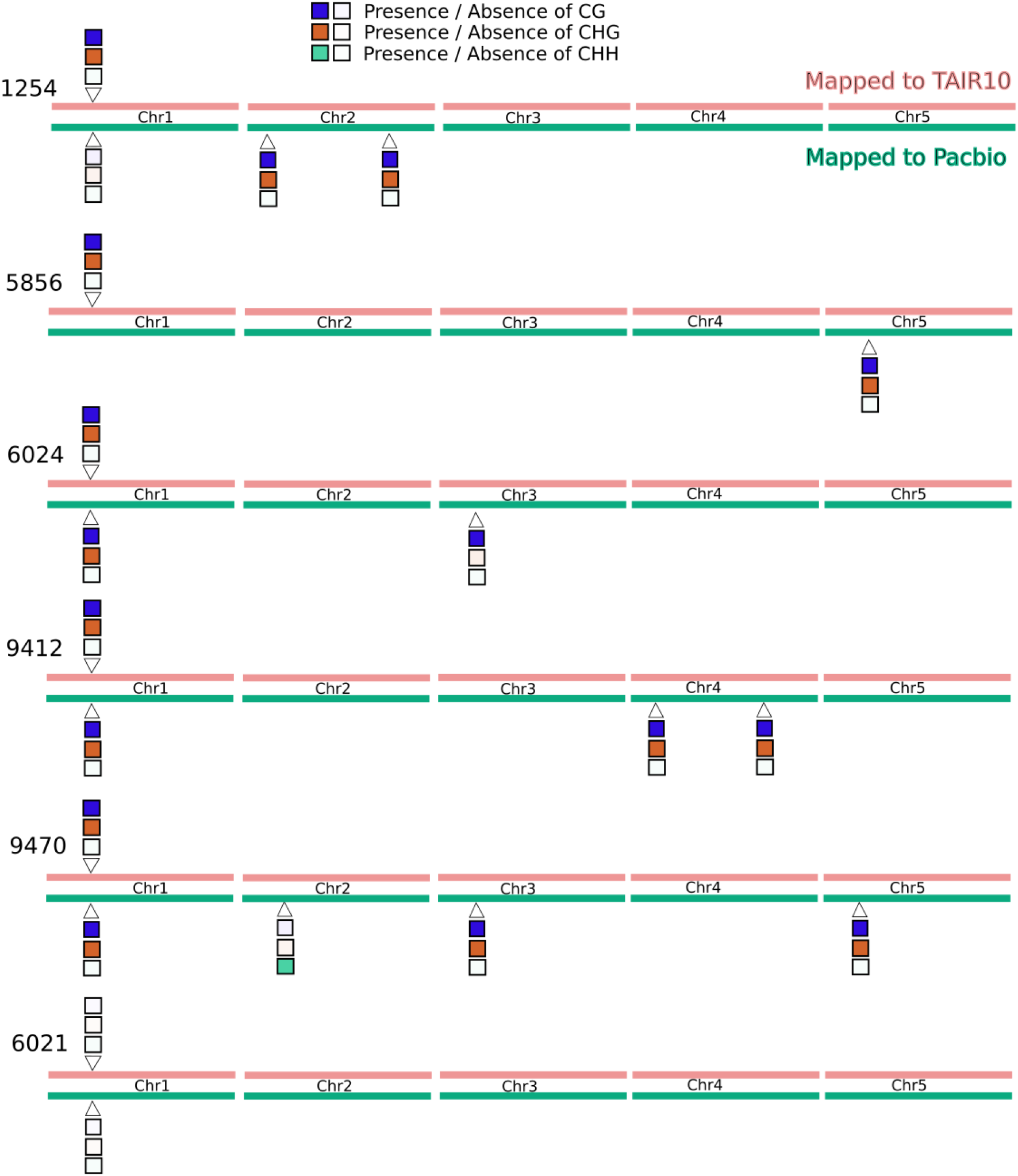
The effect of calling methylation status for AT1G30140 by mapping to a reference genome vs. the appropriate genome. Locations on the chromosomes are approximate, for illustration only.

## Discussion

A duplication can lead to pseudo-SNPs when SNPs are identified by mapping short reads to a reference genome that does not contain the duplication. Typically pseudo-SNPs have to be identified using non-Mendelian segregation patterns in families or crosses, but in inbred lines they can be identified solely by their presence. The overwhelming majority of the 3.3 million heterozygous SNPs (44% of total) identified by our SNP-calling of the 1001 Genomes Project (2016) data are likely to be pseudo-SNPs. Assuming this, we used (pseudo-)heterozygosity as a “phenotype”, and tried to map its cause, i.e. the duplication, using a simple but powerful GWAS approach. Focusing on annotated genes, we find that over 2500 (roughly 10% of total) harbor pseudo-SNPs and show evidence of duplication. Using 6 new long-read assemblies, we were able to confirm 60% of these duplications using conservative criteria (see Methods). Most of the remaining duplications are located in pericentromeric regions where SNP-calling has lower quality, and which are difficult to assemble even with long-read (**Supplemental Figure 16**).

These numbers nearly certainly underestimate the true extent of duplication, which has been known to be common in *A. thaliana* for over a decade (Cao et al. 2011; Gan et al. 2011; Schneeberger et al. 2011). While unlinked *trans-*duplications are fairly likely to give rise to pseudo-SNPs, local *cis-*duplications will only do so once sufficient time has passed for substantial sequence divergence to occur, or if they arise via non-homologous recombination in a heterozygous individual (which is less likely in *A. thaliana*). As for the GWAS approach, it lacks statistical power to detect rare duplications, and can be misled by allelic heterogeneity (due to multiple independent duplications). Finally, duplications are just a subset of structural variants, and it is therefore not surprising that other short-read approaches to detect such variants have identified many more using the 1001 Genomes data (Zmienko et al. 2020; D.-X. Liu et al. 2021; Göktay, Fulgione, and Hancock 2020).

Pseudo-SNPs is not the only problem with relying on a reference genome. Our analysis uncovered a striking example of the potential importance of the “mobileome” in shaping genome diversity (Morgante et al. 2005): we show that an annotated gene and an annotated transposon are both part of a much large mobile element, and the insertion in the reference genome is missing from most other accessions. When short reads from another accession are mapped to this “gene” using the reference genome, you are neither mapping to a gene, nor to the position you think. One possible consequence of this is incorrect methylation polymorphism calls, as we demonstrate above, but essentially any methodology that relies on mapping sequencing data to a reference genome could be affected (e.g. RNA-seq).

Time (and more independently assembled genomes) will tell how significant this problem is, but the potential for artifactual results is clearly substantial, and likely depends on the amount of recent transposon activity (Morgante et al. 2005). It is also important to realize that the artefactual nature of the 44% heterozygous SNPs was only apparent because we are working with inbred lines. Other researchers working on inbred lines have reached similar conclusions, and used various methods to eliminate them *e*.*g. Zea* (Chia et al. 2012; Lu et al. 2015; Bukowski et al. 2018) and *Brachypodium* (Stritt et al. 2021). In human genetics, SNP-calling relies heavily on family trios, but in outcrossing organisms where this is not possible, there is great cause for concern. The increasing ease and ability to sequence more and more complex genomes, such as projects associated with the 1001G+ and Tree of Life, will allow population analyses to avoid the use of a single reference genome and reveal new mechanisms of gene duplication and structural variants such as those reported here.

## Methods

### Long-read sequencing of six *A. thaliana*

We sequenced six Swedish *A. thaliana* lines that are part of the 1001 Genomes collection (1001 Genomes Consortium 2016), ecotype ids: 1254, 5856, 6021, 6024, 9412 and 9470. Plants were grown in the growth chamber at 21 C in long-day settings for 3 weeks and dark-treated for 24-48 hours before being collected. DNA was extracted from ∼20 g of frozen whole seedling material following a high molecular weight DNA extraction protocol adapted for plant tissue (Cristina Barragan et al. 2021). All six genomes were sequenced with PacBio technology, 6021 with PacBio RSII, and the rest with Sequel. Accession 9412 was sequenced twice and 6024 was additionally sequenced with Nanopore (4.1 Gbp sequenced, 376 K reads with N50 18.7 Kb). All data were used in the assemblies.

### MinION sequencing of two *A. lyrata*

We sequenced two North American *A. lyrata* accessions, 11B02 and 11B21. Both individuals come from the 11B population of *A. lyrata*, which is self-compatible and located in Missouri (Griffin and Willi 2014) (GPS coordinates 38° 28’ 07.1’’ N; 90° 42’ 34.3’’ W). Plants were bulked for 1 generation in the lab and DNA was extracted from ∼20g of 3-week old seedlings, grown at 21°C and dark treated for 3 days prior to tissue collection. DNA was extracted using a modified protocol for high molecular-weight DNA extraction from plant tissue. DNA quality was assessed with a Qubit fluorometer and a Nanodrop analysis. We used a Spot-ON Flow Cell FLO-MIN106D R9 Version with a ligation sequencing kit SQK-LSK109. Bases were called using guppy version 3.2.6 (https://nanoporetech.com/community). The final output of MinION sequencing for 11B02 was 13,67 Gbp in 763,800 reads and an N50 of 31,15 Kb. The final output of MinION sequencing for 11B21 was 17.55 Gb, 1.11 M reads with an N50 of 33.26 Kb.

### Genome assembly, polishing and scaffolding

The six *A. thaliana* genomes (ecotype ids 1254, 5856, 6021, 6024, 9412 and 9470) were assembled using Canu (v 1.7.1) (Koren et al. 2017) with default settings, except for genomeSize. Previous estimates of flow cytometry were used for this parameter (Long et al. 2013) when available or 170m was used. The values were 170m, 178m, 135m, 170m, 170m and 170m, respectively. The assemblies were corrected with two rounds of arrow (PacBio’s SMRT Link software release 5.0.0.6792) and one of Pilon (Walker et al. 2014). For arrow, the respective long reads were used and for Pilon, the 1001 Genomes DNA sequencing data, plus PCR-free Illumina 150bp data that was generated for accessions 6024 and 9412; lines 5856, 6021, 9470 had available PCR-free data (250bp reads generated by David Jaffe, Broad Institute). This resulted in 125.6Mb, 124.3Mb, 124.5Mb, 124.7Mb, 127.1Mb and 128Mb assembled bases, respectively; contained in 99, 436, 178, 99, 109 and 124 contigs, respectively. The polished contigs were ordered and scaffolded with respect to the Col-0 reference genome, using RaGOO (Alonge et al. 2019).

We assembled the genome of the two *A. lyrata* accessions 11B02 and 11B21 using Canu (Koren et al. 2017) (v 1.8) with default settings and a genome size set to 200Mb. The genomes of 11B02 and 11B21 were contained in 498 and 265 contigs, respectively. The contig assemblies were polished using Racon (Vaser et al. 2017) (v 1.4) and ONT long reads were mapped using nglmr (Sedlazeck et al. 2018) (v 0.2.7). Assemblies were further polished by mapping PCR-free Illumina 150bp short reads (∼100X for 11B02 and ∼88X for 11B21) to the long-read corrected assemblies. Short-read correction of assembly errors was carried out using Pilon (Walker et al. 2014) (v1.23). Contigs were scaffolded into pseudo-chromosomes using RaGOO (Alonge et al. 2019) and by using the error corrected long reads from Canu and the *A. lyrata* reference genome (Hu et al. 2011) and the *A. arenosa* subgenome of *A. suecica (Burns et al. 2021)* as a guide followed by manual inspection of regions. The assembly size for 11B02 was 213Mb and 11B21 was 202Mb. Genome size was estimated using findGSE (Sun et al. 2018) with a resulting estimated genome size of ∼256Mb for 11B02 and ∼237Mb for 11B21.

### Heterozygous SNPs calling / extraction

We downloaded short-read data for 1,057 accessions from the 1001 Genomes Project (1001 Genomes Consortium 2016). Raw paired-end reads were processed with cutadapt (v1.9) (Martin 2011) to remove 3’ adapters, and to trim 5’-ends with quality 15 and 3’-ends with quality 10 or N-endings. All reads were aligned to the *A. thaliana* TAIR10 reference genome (Arabidopsis Genome Initiative 2000) with BWA-MEM (v0.7.8) (H. Li 2013), and both Samtools (v0.1.18) and Sambamba (v0.6.3) were used for various file format conversions, sorting and indexing (H. Li et al. 2009; Tarasov et al. 2015), while duplicated reads where by marked by Markduplicates from Picard (v1.101; http://broadinstitute.github.io/picard/). Further steps were carried out with GATK (v3.4) functions (Van der Auwera et al. 2013; DePristo et al. 2011). Local realignment around indels were done with ‘RealignerTargetCreator’ and ‘IndelRealigner’, and base recalibration with ‘BaseRecalibrator’ by providing known indels and SNPS from The 1001 Genomes Consortium (1001 Genomes Consortium 2016). Genetic variants were called with ‘HaplotypeCaller’ in individual samples followed by joint genotyping of a single cohort with ‘GenotypeGVCFs’. An initial SNP filtering was done following the variant quality score recalibration (VQSR) protocol. Briefly, a subset of ∼181,000 high quality SNPs from the RegMap panel (Horton et al. 2012) were used as the training set for VariantRecalibrator with a priori probability of 15 and four maximum Gaussian distributions. Finally, only bi-allelic SNPs within at a sensitivity tranche level of 99.5 were kept, for a total of 7,311,237 SNPs.

### Heterozygous stretches analysis

From the VCF, Plink was used to generate .ped and .map files. (http://pngu.mgh.harvard.edu/purcell/plink/) (Purcell et al. 2007). To detect and characterize the stretches of heterozygosity the package “detectRUNS” in R was then used. (https://github.com/bioinformatics-ptp/detectRUNS/tree/master/detectRUNS). We used the function slidingRuns.run with the following parameters: WindowSize=10, threshold=0.05, RoHet=True, minDensity=1/100, rest as default.

### SNP filtering

From the raw VCF files SNP positions containing heterozygous labels were extracted using GATK VariantFiltration. From the 3.3 millions of heterozygous SNPs extracted, two filtering steps were then applied. Only SNPs with a frequency of at least 5% of the population and located in TAIR10-annotated coding regions were kept. After those filtering steps a core set of 26647 SNPs were retained for further analysis (**see Supplemental Figure 3**). Gene names and features containing those pseudo-SNPs were extracted from the TAIR10 annotation.

### GWAS

The presence and absence of pseudo-heterozygosity (coded as 1 and 0 respectively) was used as a phenotype to run GWAS. As a genotype the matrix published by the 1001 Genomes Consortium containing 10 million SNPs was been used (1001 Genomes Consortium 2016). To run all the GWAS, the pygwas package (https://github.com/timeu/PyGWAS) with the amm (accelerated mixed model) option was used. The raw output containing all SNPs was filtered, removing all SNPs with a minor allele frequency below 0.05 and/or a −log10(p-value) below 4. For each GWAS performed, the p-value as well as the position was used to call the peaks using the Fourier transform function in R (filterFFT), combined with the peak detection function (peakDetection), from the package NucleR 3.13, to automatically retrieve the position of each peak across the genome. From each peak, the highest SNPs within a region of +/- 10kb around the peak center were used (see the example in **Supplemental Figure 17)**. Using all 26647 SNPs, a summary table was generated with each pseudo-heterozygous SNP and each GWAS peak detected (**Supplemental Data**). This matrix was then used to generate **Figure 2C**, applying thresholds of −log10(p-value) of 20 and minor allele frequency of 0.1.

### Confirmation of GWAS results

To confirm the detected duplications, a combination of BLAST and synteny was used on the denovo-assembled genome. Only the insertions that segregate in the 6 new genomes were used (398). For each gene, the corresponding sequence from the TAIR10 annotation was located in the target genome using BLAST (**see Supplemental Figure 5**). A threshold of 70% sequence identity as well as 70% of the initial sequence length was used. The presence of a match within 20kb of the predicted peak position was interpreted as confirmation.

### Gene ontology

Out of the 2570 genes detected to be duplicated, 2396 have a gene ontology annotation. PLAZA.4 (Van Bel et al. 2018) was used to perform a gene enrichment analysis using the full genome as background. Data were then retrieved and plotted using R.

### Coverage and Methylation analysis

Bisulfite reads for the accessions were taken from 1001 methylomes (Kawakatsu et al. 2016). Reads were mapped to PacBio genomes using an nf-core pipeline (https://github.com/rbpisupati/methylseq). We filtered for cytosines with a minimum depth of 3. They methylation levels were calculated either on the gene-body or on 200bp windows using custom python scripts following guidelines from Schultz et al. (2012). Weighted methylation levels were used, i.e. if there are three cytosines with a depth of t1, t2 and t3 and number of methylated reads are c1, c2 and c3, the methylation level was calculated as (c1+c2+c3)/(t1+t2+t3). We called a gene “differentially methylated” if the difference in weighted methylation level was more than 0.05 for CG and 0.03 for CHG.

The sequencing coverage for each accession was extracted using the function bamCoverage (windows size of 50bp) from the program DeepTools (Ramírez et al. 2016). The Bigwig files generated were then processed in R using the package rtracklayer. No correlation between the mean sequencing coverage and the number of pseudo-SNPs detected was observed (**Supplemental Figure 18**).

### Multiple sequence alignment

For each insertion of the AT1G20390-AT1G20400 (Transposon+gene) fragment, a fasta file including 2kb on each side of the fragment was extracted from each genome, using the getfasta function from bedtools (Quinlan and Hall 2010). Multiple alignment was performed using KALIGN (Madeira et al. 2019). Visualization and comparison was done using Jalview 2 (Waterhouse et al. 2009).

### Structural variation analysis

To control the structure of the region around duplicated genes, the sequence from 3 genes upstream and downstream of the gene of interest was extracted. Each sequence was then BLAST to each of the genomes and the position of each BLAST result was retrieved. NCBI BLAST (Altschul et al. 1990) was used with a percentage of identity threshold of 70% and all other parameters as default. From each blast results fragments with at least 50% of the input sequence length have been selected and plotted using R.

### Frequency of the insertions in the 1001 Genomes dataset

The same sequences used for the multiple alignment were used to confirm presence or absence of each insertion in the 1001 Genomes dataset. We used each of those sequences as reference to map short reads using minimap 2 (H. Li 2018). For each insertion, only paired-end reads having both members of the pair mapping to the region were retained. An insertion was considered present in an accession if at least 3 pairs of reads spanned the insertion border **(see Supplemental Figure 11)**.

### Multiple species comparison

We used the *Capsella rubella* and *A*.*arenosa* genomes (Slotte et al. 2013; Burns et al. 2021) to search for the new Transposon+gene element, just like in the *A. thaliana* genomes. For *A. arenosa* we used the subgenome of *A. suecica*. We located the transposon+gene fragments, extracted from the TAIR10 annotation, using NCBI BLAST as above. For *A*.*lyrata* two newly assembled genomes were assembled using MinION sequencing.

## Additional files

Additional file 1.txt

Methylation value per gene of all accessions mapped to the reference genome

CG and CHG weighted average per genes of the 6 accessions analyzed. Row names correspond to the gene ID and column name to the CG and CHG for each accession.

Additional file 2-8.csv

Methylation value per gene of all accessions mapped to the corresponding genome.

CG and CHG weighted average per genes of the 6 accessions analyzed. Row names correspond to the gene ID. (the “_” corresponds to the multiple copies detected). The column name to the CG and CHG for each accession.

## Acknowledgment

We thank numerous people on Twitter for providing feedback on the bioRxiv version.

## Authors’ contributions

BJ and MN developed the project. BJ performed all analyses. LMS and RB assembled the *A*.*thaliana* and *A*.*lyrata* genomes, respectively. FR generated the SNP matrix. RP performed the methylation analyses. BJ and MN wrote the manuscript, with input from all authors.

## Availability of data and materials

All genome assemblies and raw reads were deposited under the BioProject ID: PRJNA779205. Link of the genome files for the reviewers: https://dataview.ncbi.nlm.nih.gov/object/PRJNA779205?reviewer=gduvs00c97i3bd5he06gs25oos

Scripts used are available under Github link: https://github.com/benjj212/duplication-paper.git. The full GWAS matrix is available at https://doi.org/10.5281/zenodo.5702395

## Ethics approval and consent to participate

Not applicable.

## References

1001 Genomes Consortium. 2016. “1,135 Genomes Reveal the Global Pattern of Polymorphism in Arabidopsis Thaliana.” Cell 166 (2): 481–91.

Alkan, Can, Bradley P. Coe, and Evan E. Eichler. 2011. “Genome Structural Variation Discovery and Genotyping.” Nature Reviews. Genetics 12 (5): 363–76.

Alonge, Michael, Sebastian Soyk, Srividya Ramakrishnan, Xingang Wang, Sara Goodwin, Fritz J. Sedlazeck, Zachary B. Lippman, and Michael C. Schatz. 2019. “RaGOO: Fast and Accurate Reference-Guided Scaffolding of Draft Genomes.” Genome Biology 20 (1): 224.

Alonge, Michael, Xingang Wang, Matthias Benoit, Sebastian Soyk, Lara Pereira, Lei Zhang, Hamsini Suresh, et al. 2020. “Major Impacts of Widespread Structural Variation on Gene Expression and Crop Improvement in Tomato.” Cell. https://doi.org/10.1016/j.cell.2020.05.021.

Altschul, S. F., W. Gish, W. Miller, E. W. Myers, and D. J. Lipman. 1990. “Basic Local Alignment Search Tool.” Journal of Molecular Biology 215 (3): 403–10.

Arabidopsis Genome Initiative. 2000. “Analysis of the Genome Sequence of the Flowering Plant Arabidopsis Thaliana.” Nature 408 (6814): 796–815.

Bukowski, Robert, Xiaosen Guo, Yanli Lu, Cheng Zou, Bing He, Zhengqin Rong, Bo Wang, et al. 2018. “Construction of the Third-Generation Zea Mays Haplotype Map.” GigaScience 7 (4): 1–12.

Burns, Robin, Terezie Mandáková, Joanna Gunis, Luz Mayela Soto-Jiménez, Chang Liu, Martin A. Lysak, Polina Yu Novikova, and Magnus Nordborg. 2021. “Gradual Evolution of Allopolyploidy in Arabidopsis Suecica.” Nature Ecology & Evolution 5 (10): 1367–81.

Cao, Jun, Korbinian Schneeberger, Stephan Ossowski, Torsten Günther, Sebastian Bender, Joffrey Fitz, Daniel Koenig, et al. 2011. “Whole-Genome Sequencing of Multiple Arabidopsis Thaliana Populations.” Nature Genetics 43 (10): 956–63.

Carter, Nigel P. 2007. “Methods and Strategies for Analyzing Copy Number Variation Using DNA Microarrays.” Nature Genetics 39 (7 Suppl): S16–21.

Chia, Jer-Ming, Chi Song, Peter J. Bradbury, Denise Costich, Natalia de Leon, John Doebley, Robert J. Elshire, et al. 2012. “Maize HapMap2 Identifies Extant Variation from a Genome in Flux.” Nature Genetics 44 (7): 803–7.

Cristina Barragan, A., Maximilian Collenberg, Rebecca Schwab, Merijn Kerstens, Ilja Bezrukov, Felix Bemm, Doubravka Požárová, Filip Kolář, and Detlef Weigel. 2021. “Homozygosity at Its Limit: Inbreeding Depression in Wild Arabidopsis Arenosa Populations.” bioRxiv. https://doi.org/10.1101/2021.01.24.427284.

DePristo, Mark A., Eric Banks, Ryan Poplin, Kiran V. Garimella, Jared R. Maguire, Christopher Hartl, Anthony A. Philippakis, et al. 2011. “A Framework for Variation Discovery and Genotyping Using next-Generation DNA Sequencing Data.” Nature Genetics 43 (5): 491–98.

Gan, Xiangchao, Oliver Stegle, Jonas Behr, Joshua G. Steffen, Philipp Drewe, Katie L. Hildebrand, Rune Lyngsoe, et al. 2011. “Multiple Reference Genomes and Transcriptomes for Arabidopsis Thaliana.” Nature 477 (7365): 419–23.

Göktay, Mehmet, Andrea Fulgione, and Angela M. Hancock. 2020. “A New Catalogue of Structural Variants in 1301 A. Thaliana Lines from Africa, Eurasia and North America Reveals a Signature of Balancing at Defense Response Genes.” Molecular Biology and Evolution, November. https://doi.org/10.1093/molbev/msaa309.

Gonzalez, Enrique, Hemant Kulkarni, Hector Bolivar, Andrea Mangano, Racquel Sanchez, Gabriel Catano, Robert J. Nibbs, et al. 2005. “The Influence of CCL3L1 Gene-Containing Segmental Duplications on HIV-1/AIDS Susceptibility.” Science 307 (5714): 1434–40.

Griffin, P. C., and Y. Willi. 2014. “Evolutionary Shifts to Self-Fertilisation Restricted to Geographic Range Margins in North American Arabidopsis Lyrata.” Ecology Letters 17 (4): 484–90.

Handsaker, Robert E., Joshua M. Korn, James Nemesh, and Steven A. McCarroll. 2011. “Discovery and Genotyping of Genome Structural Polymorphism by Sequencing on a Population Scale.” Nature Genetics 43 (3): 269–76.

Horton, Matthew W., Angela M. Hancock, Yu S. Huang, Christopher Toomajian, Susanna Atwell, Adam Auton, N. Wayan Muliyati, et al. 2012. “Genome-Wide Patterns of Genetic Variation in Worldwide Arabidopsis Thaliana Accessions from the RegMap Panel.” Nature Genetics 44 (2): 212–16.

Hufford, Matthew B., Arun S. Seetharam, Margaret R. Woodhouse, Kapeel M. Chougule, Shujun Ou, Jianing Liu, William A. Ricci, et al. 2021. “De Novo Assembly, Annotation, and Comparative Analysis of 26 Diverse Maize Genomes.” Cold Spring Harbor Laboratory. https://doi.org/10.1101/2021.01.14.426684.

Hurles, Matthew. 2002. “Are 100,000 ‘SNPs’ Useless?” Science.

Hu, Tina T., Pedro Pattyn, Erica G. Bakker, Jun Cao, Jan-Fang Cheng, Richard M. Clark, Noah Fahlgren, et al. 2011. “The Arabidopsis Lyrata Genome Sequence and the Basis of Rapid Genome Size Change.” Nature Genetics 43 (5): 476–81.

Jiao, Wen-Biao, and Korbinian Schneeberger. 2019. “Chromosome-Level Assemblies of Multiple Arabidopsis Thaliana Accessions Reveal Hotspots of Genomic Rearrangements.” bioRxiv. https://doi.org/10.1101/738880.

Kawakatsu, Taiji, Shao-Shan Carol Huang, Florian Jupe, Eriko Sasaki, Robert J. Schmitz, Mark A. Urich, Rosa Castanon, et al. 2016. “Epigenomic Diversity in a Global Collection of Arabidopsis Thaliana Accessions.” Cell 166 (2): 492–505.

Koren, Sergey, Brian P. Walenz, Konstantin Berlin, Jason R. Miller, Nicholas H. Bergman, and Adam M. Phillippy. 2017. “Canu: Scalable and Accurate Long-Read Assembly via Adaptive K-Mer Weighting and Repeat Separation.” Genome Research 27 (5): 722–36.

Li, Changsheng, Xiaoli Xiang, Yongcai Huang, Yong Zhou, Dong An, Jiaqiang Dong, Chenxi Zhao, et al. 2020. “Long-Read Sequencing Reveals Genomic Structural Variations That Underlie Creation of Quality Protein Maize.” Nature Communications 11 (1): 17.

Li, Heng. 2013. “Aligning Sequence Reads, Clone Sequences and Assembly Contigs with BWA-MEM.” arXiv [q-bio.GN]. arXiv. http://arxiv.org/abs/1303.3997.

Li, Heng. 2018. “Minimap2: Pairwise Alignment for Nucleotide Sequences.” Bioinformatics 34 (18): 3094–3100.

Li, Heng, Bob Handsaker, Alec Wysoker, Tim Fennell, Jue Ruan, Nils Homer, Gabor Marth, Goncalo Abecasis, Richard Durbin, and 1000 Genome Project Data Processing Subgroup. 2009. “The Sequence Alignment/Map Format and SAMtools.” Bioinformatics 25 (16): 2078–79.

Lin, Ke, Ningwen Zhang, Edouard I. Severing, Harm Nijveen, Feng Cheng, Richard G. F. Visser, Xiaowu Wang, Dick de Ridder, and Guusje Bonnema. 2014. “Beyond Genomic Variation - Comparison and Functional Annotation of Three Brassica Rapagenomes: A Turnip, a Rapid Cycling and a Chinese Cabbage.” BMC Genomics 15 (1): 250.

Lisch, Damon. 2013. “How Important Are Transposons for Plant Evolution?” Nature Reviews. Genetics 14 (1): 49–61.

Liu, Dong-Xu, Ramesh Rajaby, Lu-Lu Wei, Lei Zhang, Zhi-Quan Yang, Qing-Yong Yang, and Wing-Kin Sung. 2021. “Calling Large Indels in 1047 Arabidopsis with IndelEnsembler.” Nucleic Acids Research, October. https://doi.org/10.1093/nar/gkab904.

Liu, Yucheng, Huilong Du, Pengcheng Li, Yanting Shen, Hua Peng, Shulin Liu, Guo-An Zhou, et al. 2020. “Pan-Genome of Wild and Cultivated Soybeans.” Cell, June. https://doi.org/10.1016/j.cell.2020.05.023.

Long, Quan, Fernando A. Rabanal, Dazhe Meng, Christian D. Huber, Ashley Farlow, Alexander Platzer, Qingrun Zhang, et al. 2013. “Massive Genomic Variation and Strong Selection in Arabidopsis Thaliana Lines from Sweden.” Nature Genetics 45 (8): 884–90.

Lu, Fei, Maria C. Romay, Jeffrey C. Glaubitz, Peter J. Bradbury, Robert J. Elshire, Tianyu Wang, Yu Li, et al. 2015. “High-Resolution Genetic Mapping of Maize Pan-Genome Sequence Anchors.” Nature Communications 6 (April): 6914.

Madeira, Fábio, Young Mi Park, Joon Lee, Nicola Buso, Tamer Gur, Nandana Madhusoodanan, Prasad Basutkar, et al. 2019. “The EMBL-EBI Search and Sequence Analysis Tools APIs in 2019.” Nucleic Acids Research 47 (W1): W636–41.

Martin, Marcel. 2011. “Cutadapt Removes Adapter Sequences from High-Throughput Sequencing Reads.” EMBnet.journal 17 (1): 10–12.

Melquist, S., B. Luff, and J. Bender. 1999. “Arabidopsis PAI Gene Arrangements, Cytosine Methylation and Expression.” Genetics 153 (1): 401–13.

Miyahara, E., J. Pokorny, V. C. Smith, R. Baron, and E. Baron. 1998. “Color Vision in Two Observers with Highly Biased LWS/MWS Cone Ratios.” Vision Research 38 (4): 601–12.

Morgante, Michele, Stephan Brunner, Giorgio Pea, Kevin Fengler, Andrea Zuccolo, and Antoni Rafalski. 2005. “Gene Duplication and Exon Shuffling by Helitron-like Transposons Generate Intraspecies Diversity in Maize.” Nature Genetics 37 (9): 997–1002.

Perry, George H., Nathaniel J. Dominy, Katrina G. Claw, Arthur S. Lee, Heike Fiegler, Richard Redon, John Werner, et al. 2007. “Diet and the Evolution of Human Amylase Gene Copy Number Variation.” Nature Genetics 39 (10): 1256–60.

Pinosio, Sara, Stefania Giacomello, Patricia Faivre-Rampant, Gail Taylor, Veronique Jorge, Marie Christine Le Paslier, Giusi Zaina, et al. 2016. “Characterization of the Poplar Pan-Genome by Genome-Wide Identification of Structural Variation.” Molecular Biology and Evolution 33 (10): 2706–19.

Purcell, Shaun, Benjamin Neale, Kathe Todd-Brown, Lori Thomas, Manuel A. R. Ferreira, David Bender, Julian Maller, et al. 2007. “PLINK: A Tool Set for Whole-Genome Association and Population-Based Linkage Analyses.” American Journal of Human Genetics 81 (3): 559–75.

Quadrana, Leandro, Amanda Bortolini Silveira, George F. Mayhew, Chantal LeBlanc, Robert A. Martienssen, Jeffrey A. Jeddeloh, Vincent Colot, and Daniel Zilberman. 2016. “The Arabidopsis Thaliana Mobilome and Its Impact at the Species Level.” eLife 5 (June): e15716.

Quinlan, Aaron R., and Ira M. Hall. 2010. “BEDTools: A Flexible Suite of Utilities for Comparing Genomic Features.” Bioinformatics 26 (6): 841–42.

Ramírez, Fidel, Devon P. Ryan, Björn Grüning, Vivek Bhardwaj, Fabian Kilpert, Andreas S. Richter, Steffen Heyne, Friederike Dündar, and Thomas Manke. 2016. “deepTools2: A next Generation Web Server for Deep-Sequencing Data Analysis.” Nucleic Acids Research 44 (W1): W160–65.

Ranade, K., M. S. Chang, C. T. Ting, D. Pei, C. F. Hsiao, M. Olivier, R. Pesich, et al. 2001. “High-Throughput Genotyping with Single Nucleotide Polymorphisms.” Genome Research 11 (7): 1262–68.

Schneeberger, Korbinian, Stephan Ossowski, Felix Ott, Juliane D. Klein, Xi Wang, Christa Lanz, Lisa M. Smith, et al. 2011. “Reference-Guided Assembly of Four Diverse Arabidopsis Thaliana Genomes.” Proceedings of the National Academy of Sciences of the United States of America 108 (25): 10249–54.

Schultz, Matthew D., Robert J. Schmitz, and Joseph R. Ecker. 2012. “‘Leveling’ the Playing Field for Analyses of Single-Base Resolution DNA Methylomes.” Trends in Genetics: TIG 28 (12): 583–85.

Sedlazeck, Fritz J., Philipp Rescheneder, Moritz Smolka, Han Fang, Maria Nattestad, Arndt von Haeseler, and Michael C. Schatz. 2018. “Accurate Detection of Complex Structural Variations Using Single-Molecule Sequencing.” Nature Methods 15 (6): 461–68.

Shendure, Jay, and Hanlee Ji. 2008. “Next-Generation DNA Sequencing.” Nature Biotechnology 26 (10): 1135–45.

Slotte, Tanja, Khaled M. Hazzouri, J. Arvid Ågren, Daniel Koenig, Florian Maumus, Ya-Long Guo, Kim Steige, et al. 2013. “The Capsella Rubella Genome and the Genomic Consequences of Rapid Mating System Evolution.” Nature Genetics 45 (7): 831–35.

Snijders, A. M., N. Nowak, R. Segraves, S. Blackwood, N. Brown, J. Conroy, G. Hamilton, et al. 2001. “Assembly of Microarrays for Genome-Wide Measurement of DNA Copy Number.” Nature Genetics 29 (3): 263–64.

Stritt, Christoph, Elena L. Gimmi, Michele Wyler, Abdelmonaim H. Bakali, Aleksandra Skalska, Robert Hasterok, Luis A. J. Mur, Nicola Pecchioni, and Anne C. Roulin. 2021. “Migration without Interbreeding: Evolutionary History of a Highly Selfing Mediterranean Grass Inferred from Whole Genomes.” Molecular Ecology, October. https://doi.org/10.1111/mec.16207.

Sun, Hequan, Jia Ding, Mathieu Piednoël, and Korbinian Schneeberger. 2018. “findGSE: Estimating Genome Size Variation within Human and Arabidopsis Using K-Mer Frequencies.” Bioinformatics 34 (4): 550–57.

Tarasov, Artem, Albert J. Vilella, Edwin Cuppen, Isaac J. Nijman, and Pjotr Prins. 2015. “Sambamba: Fast Processing of NGS Alignment Formats.” Bioinformatics 31 (12): 2032–34.

Van Bel, Michiel, Tim Diels, Emmelien Vancaester, Lukasz Kreft, Alexander Botzki, Yves Van de Peer, Frederik Coppens, and Klaas Vandepoele. 2018. “PLAZA 4.0: An Integrative Resource for Functional, Evolutionary and Comparative Plant Genomics.” Nucleic Acids Research 46 (D1): D1190–96.

Van der Auwera, Geraldine A., Mauricio O. Carneiro, Chris Hartl, Ryan Poplin, Guillermo Del Angel, Ami Levy-Moonshine, Tadeusz Jordan, et al. 2013. “From FastQ Data to High Confidence Variant Calls: The Genome Analysis Toolkit Best Practices Pipeline.” Current Protocols in Bioinformatics / Editoral Board, Andreas D. Baxevanis … [et Al.] 11 (1110): 11.10.1–11.10.33.

Vaser, Robert, Ivan Sović, Niranjan Nagarajan, and Mile Šikić. 2017. “Fast and Accurate de Novo Genome Assembly from Long Uncorrected Reads.” Genome Research 27 (5): 737–46.

Walker, Bruce J., Thomas Abeel, Terrance Shea, Margaret Priest, Amr Abouelliel, Sharadha Sakthikumar, Christina A. Cuomo, et al. 2014. “Pilon: An Integrated Tool for Comprehensive Microbial Variant Detection and Genome Assembly Improvement.” PloS One 9 (11): e112963.

Waterhouse, Andrew M., James B. Procter, David M. A. Martin, Michèle Clamp, and Geoffrey J. Barton. 2009. “Jalview Version 2--a Multiple Sequence Alignment Editor and Analysis Workbench.” Bioinformatics 25 (9): 1189–91.

Woodhouse, Margaret R., Brent Pedersen, and Michael Freeling. 2010. “Transposed Genes in Arabidopsis Are Often Associated with Flanking Repeats.” PLoS Genetics 6 (5): e1000949.

Yao, Wen, Guangwei Li, Hu Zhao, Gongwei Wang, Xingming Lian, and Weibo Xie. 2015. “Exploring the Rice Dispensable Genome Using a Metagenome-like Assembly Strategy.” Genome Biology 16 (September): 187.

Zhao, Min, Qingguo Wang, Quan Wang, Peilin Jia, and Zhongming Zhao. 2013. “Computational Tools for Copy Number Variation (CNV) Detection Using next-Generation Sequencing Data: Features and Perspectives.” BMC Bioinformatics 14 Suppl 11 (September): S1.

Zhou, Yong, Dmytro Chebotarov, Dave Kudrna, Victor Llaca, Seunghee Lee, Shanmugam Rajasekar, Nahed Mohammed, et al. 2020. “A Platinum Standard Pan-Genome Resource That Represents the Population Structure of Asian Rice.” Scientific Data 7 (1): 113.

Zmienko, Agnieszka, Malgorzata Marszalek-Zenczak, Pawel Wojciechowski, Anna Samelak-Czajka, Magdalena Luczak, Piotr Kozlowski, Wojciech M. Karlowski, and Marek Figlerowicz. 2020. “AthCNV: A Map of DNA Copy Number Variations in the Arabidopsis Genome.” The Plant Cell 32 (6): 1797–1819.

