## Supplemental Figures for "Extensive gene duplication in Arabidopsis revealed by pseudo-heterozygosity"

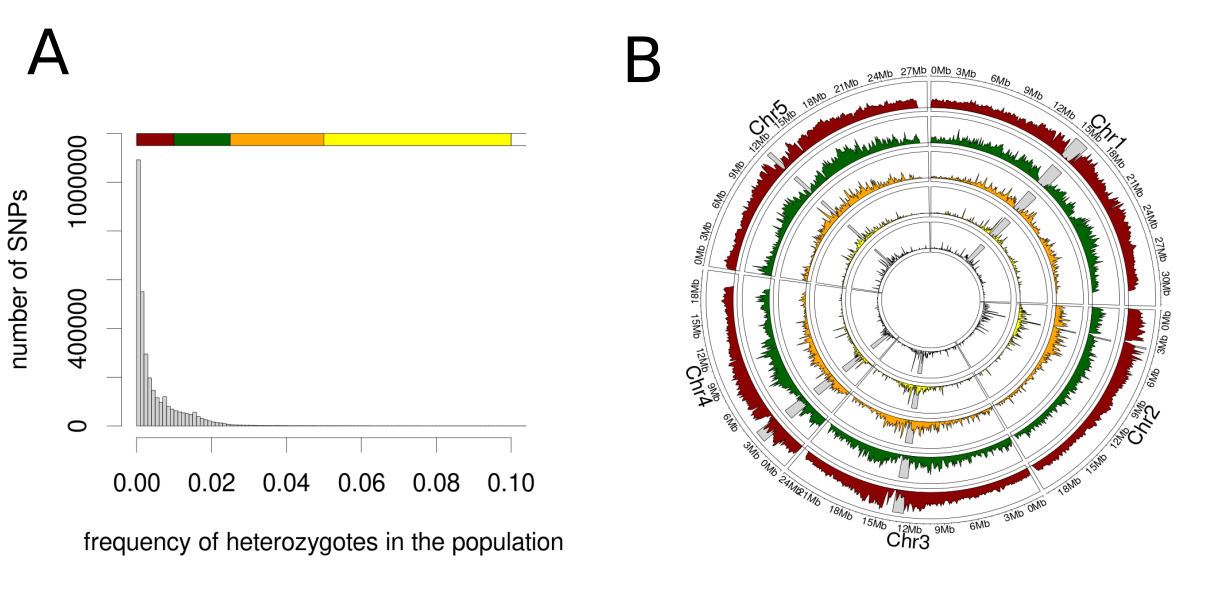

**Supplemental Figure 1:** Distribution of heterozygosity across individuals and the genome.

**(A)** Frequency distribution of the heterozygosity at individual SNPs in 1135 accessions. **(B)** Circular plot of the density of pseudo-heterozygous SNPs at different frequencies (indicated by colors corresponding to the bar in plot A)

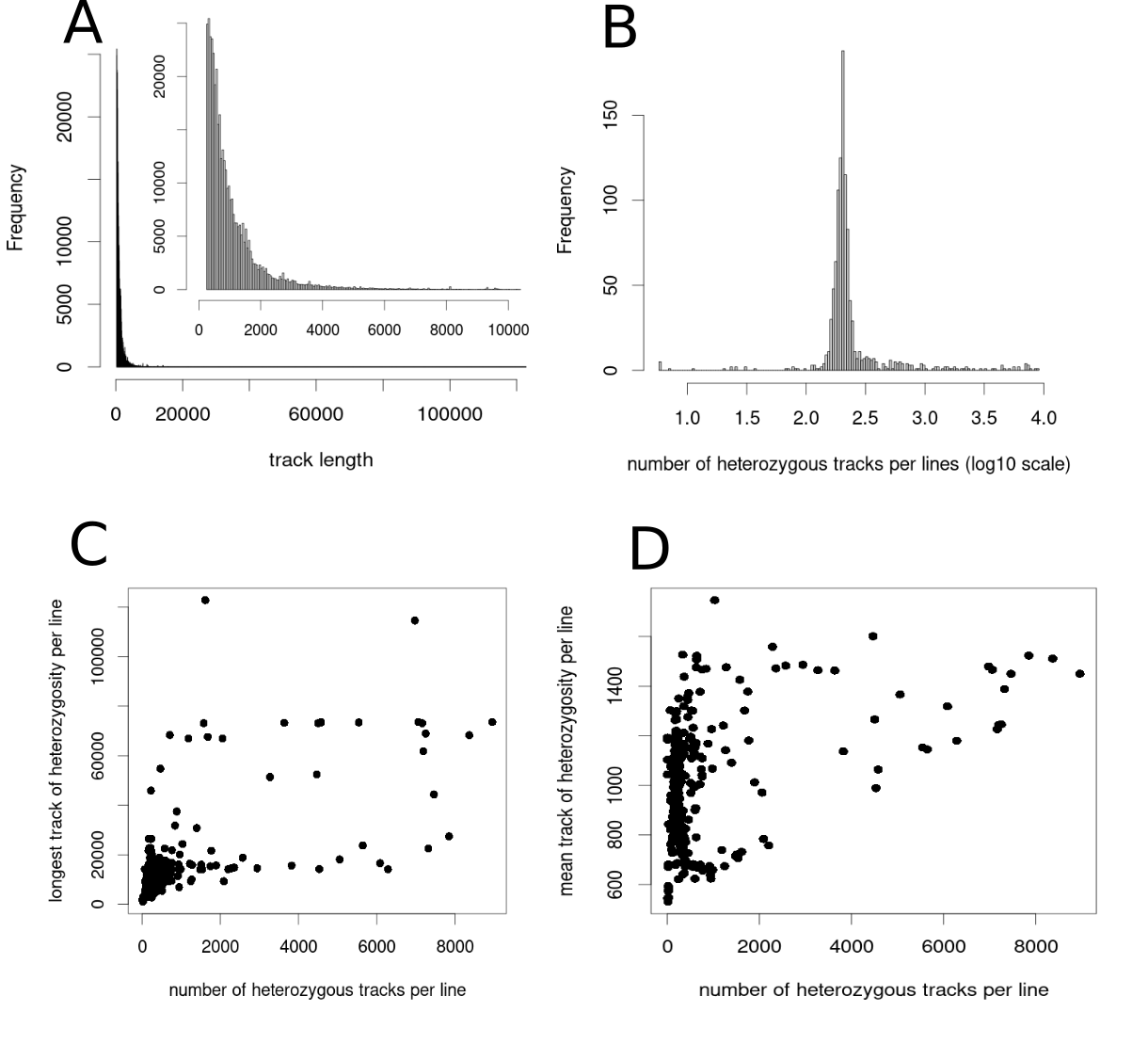

**Supplemental Figure 2:** Tract-length distribution. **(A)** The length distribution of heterozygous tracts across the 1001 genomes. **(B)** The distribution of the number of heterozygous tracts per genome. **(C)** The relationship between the maximum length of a heterozygous tract per line and the number of tracts per line. **(D)** The relationship between the mean length of heterozygous tracts per line and the number of tracts per line.

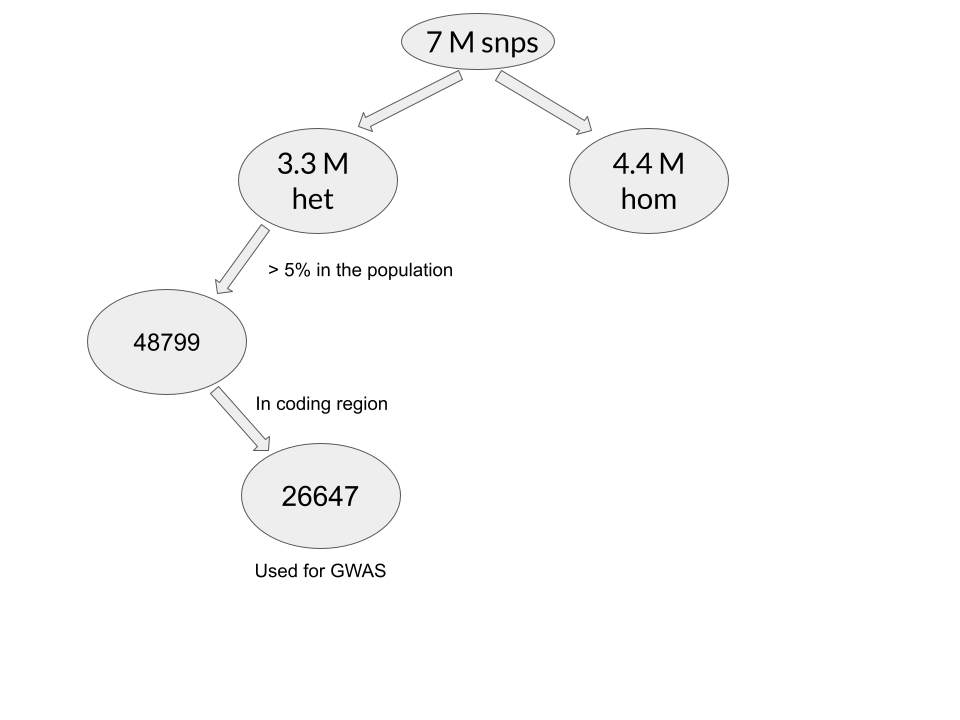

**Supplemental figure 3:** SNP filtering scheme. The SNPs matrix we started with contains 7 million SNPs. Of those, 3.3 million were detected as heterozygous in at least one line. We selected the 48,799 SNPs that we called heterozygous in at least 5 % of the lines, and focused on the 26,647 found in coding regions (according to the TAIR10 annotation).

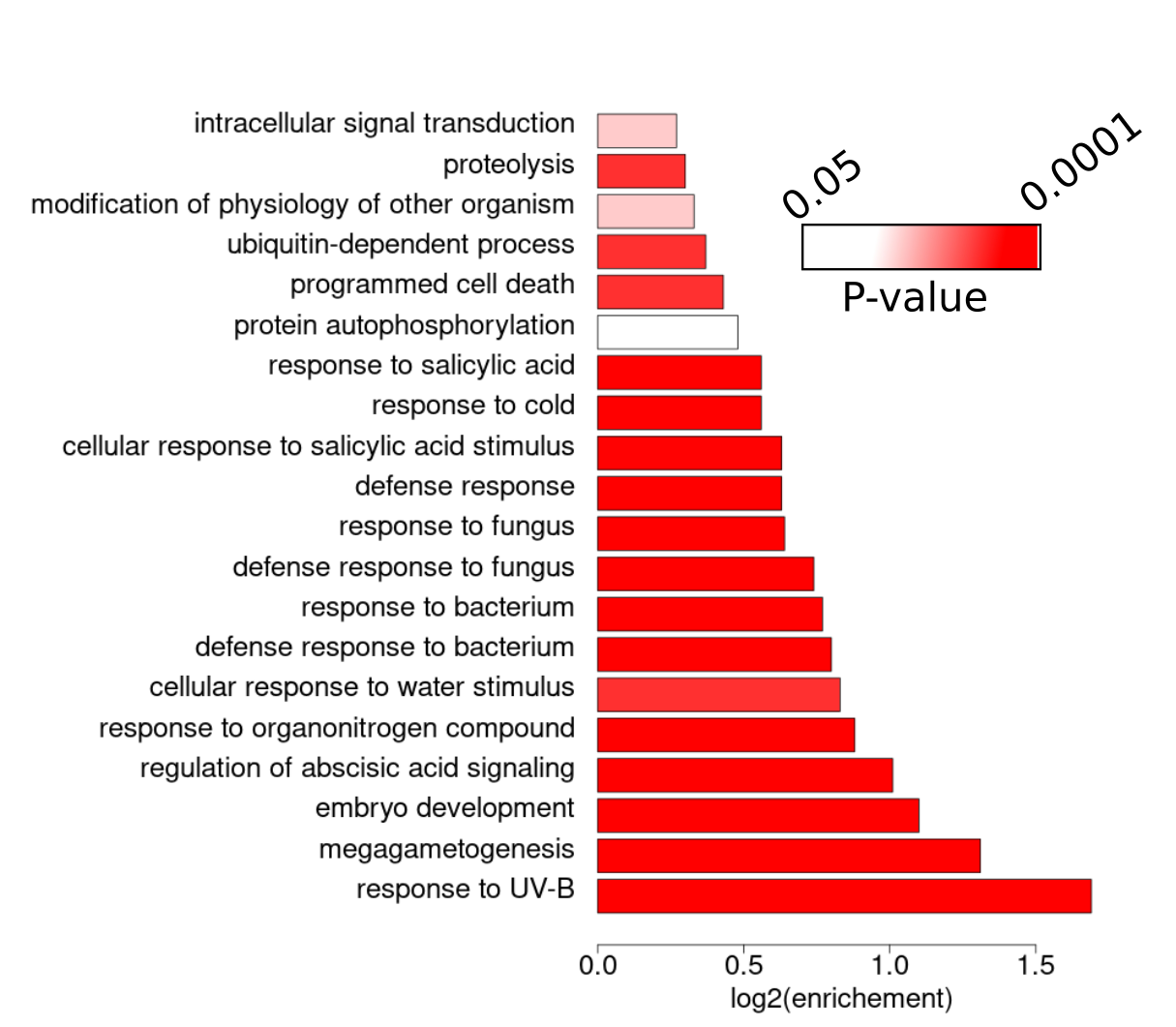

**Supplemental Figure 4:** Gene ontology analysis of the putatively duplicated genes.

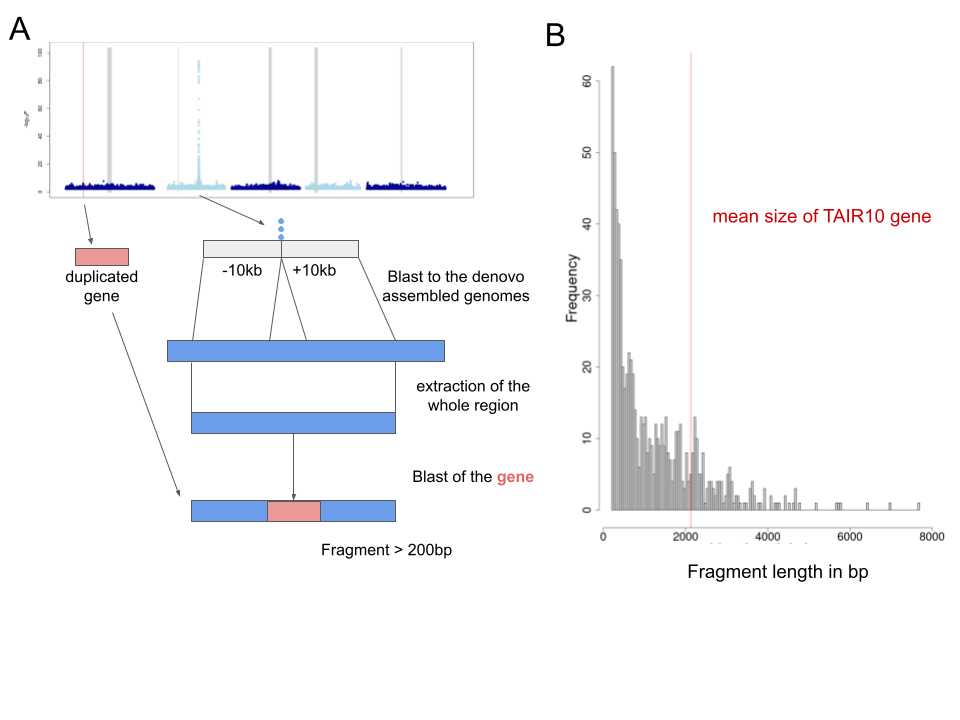

**Supplemental Figure 5:** Pipeline to confirm GWAS peaks. **(A)** For each peak detected, the flanking region (+/- 10kb) in the reference genome was extracted and located in each of the newly assembled genomes using BLAST (Altschul et al. 1990). The longest possible matches (minimum size 2kb) were extracted. The gene of interest (from TAIR10) was then aligned to this fragment using BLAST, and the result were analyzed as described in Methods. **(B)** Frequency of the initial matches found across the 6 Pacbio genomes. Only fragments longer than 200 bp were considered.

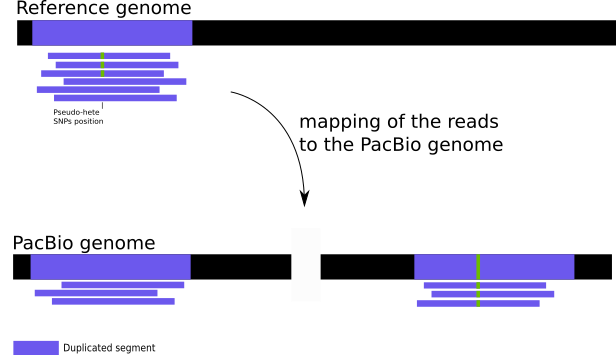

**Supplemental Figure 6:** Mapping reads tagging singleton pseudo-SNP. First, all reads overlapping the position of a specific pseudo-SNP were extracted based on mapping to the reference genome (TAIR10). This set of reads were then re-mapped to the appropriate Pacbio genome. Reads mapping to multiple regions indicate the presence of a duplicated segment. A decrease in coverage compared to the mapping to the reference genome is also a confirmation that reads map at different positions. An example is presented in **Supplement Figure 8**.

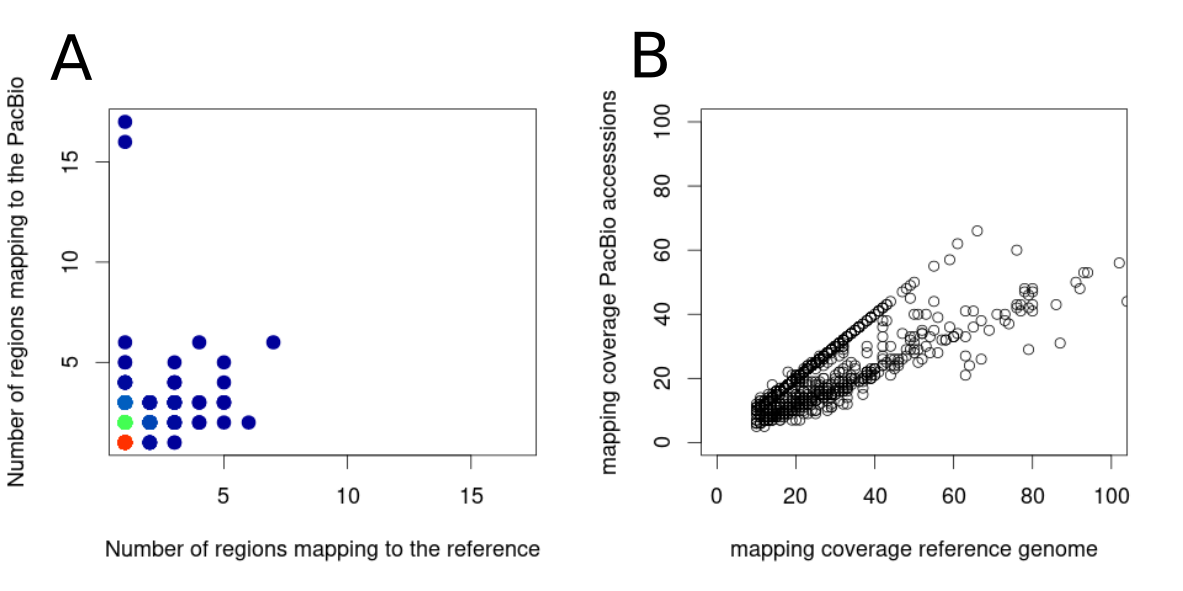

**Supplemental Figure 7:** Comparison of mapping coverage between reference genome and PacBio genomes for all regions surrounding pseudo-heterozygous doubleton positions. **(A)** Scatterplot comparing the number region found when mapping reads extracted from doubleton to the reference or to the corresponding newly assembled genome. Colors represent the density of dots. (blue = low density and red = high density). **(B)** Scatterplot comparing the coverage at doubleton pseudo-SNPs when mapped to the reference genome and when mapped to the appropriate PacBio genomes.

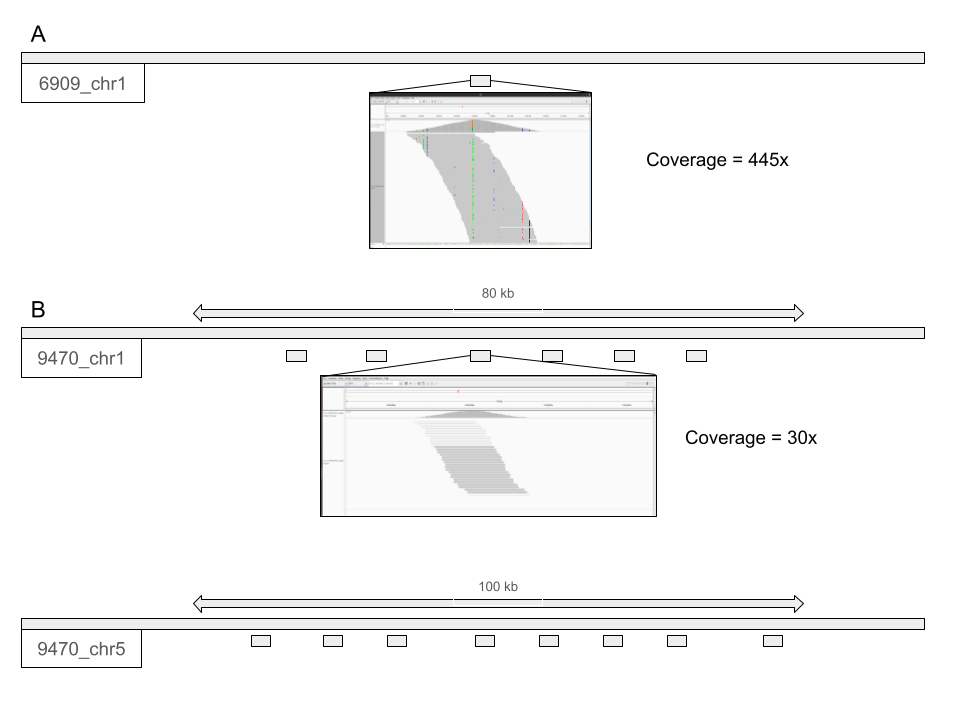

**Supplemental Figure 8:** Mapping of reads overlapping singletons (example). **(A)** Genome browser view of the region surrounding a selected pseudo-SNP. This mapping includes only the reads overlapping the pseudo-SNP position. **(B)** Mapping positions on a newly assembled genome (accession 9470). The mapping includes all the reads extracted from A.

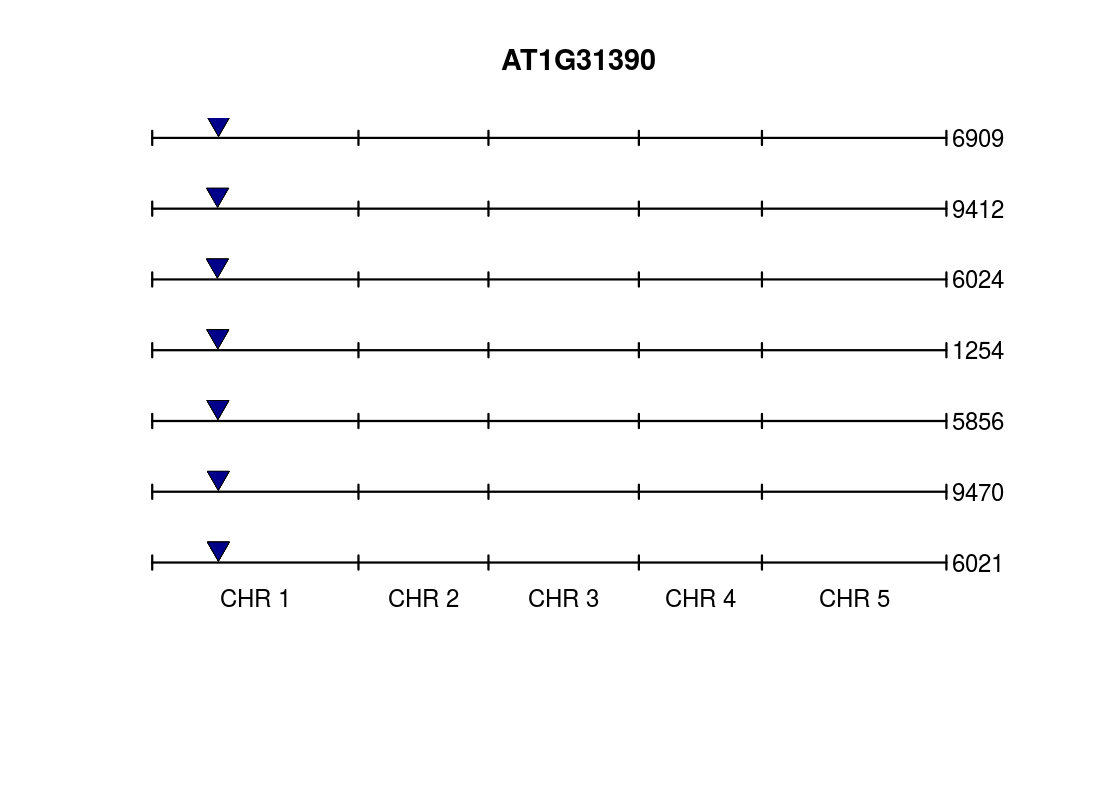

**Supplemental Figure 9:** Position of AT1G31390 in the reference (accession 6909) and each of the six newly assembled genomes. BLAST-thresholds of 70% identity were used, and only fragments of length greater than 50% of the original gene length are shown.

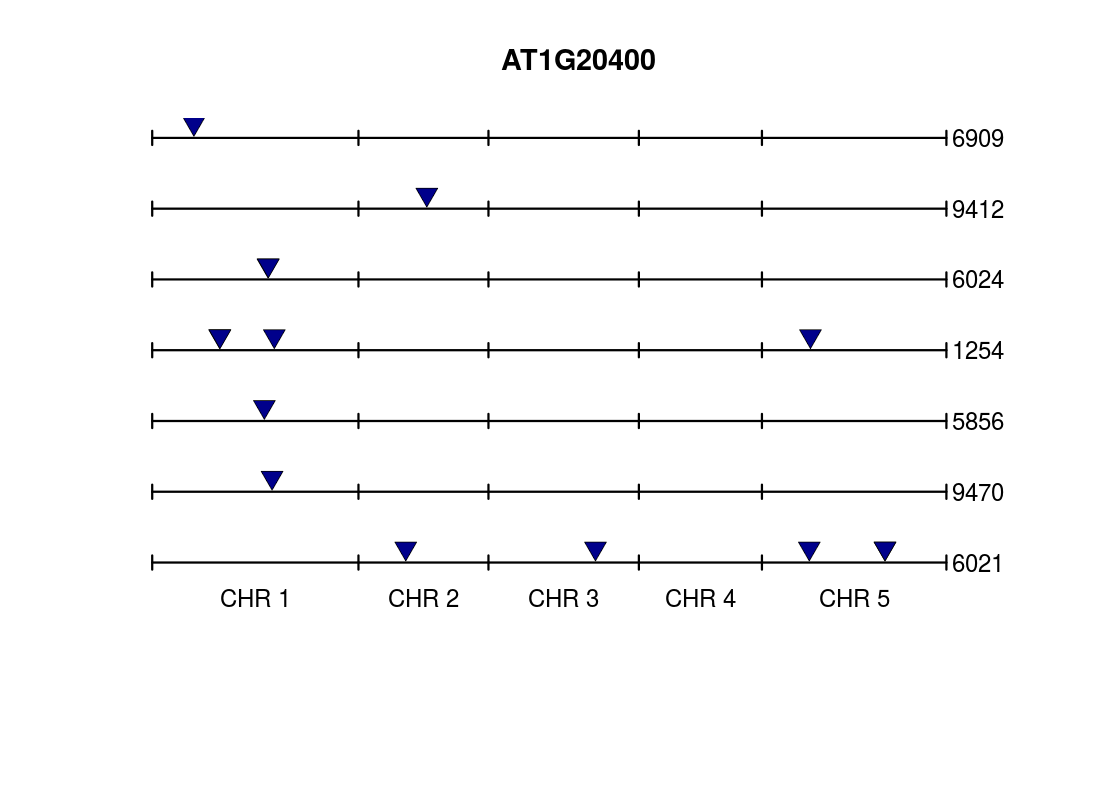

**Supplemental Figure 10:** Position of AT1G20400 in the reference (accession 6909) and each of the six newly assembled genomes. Cf. **Supplemental Figure 9**.

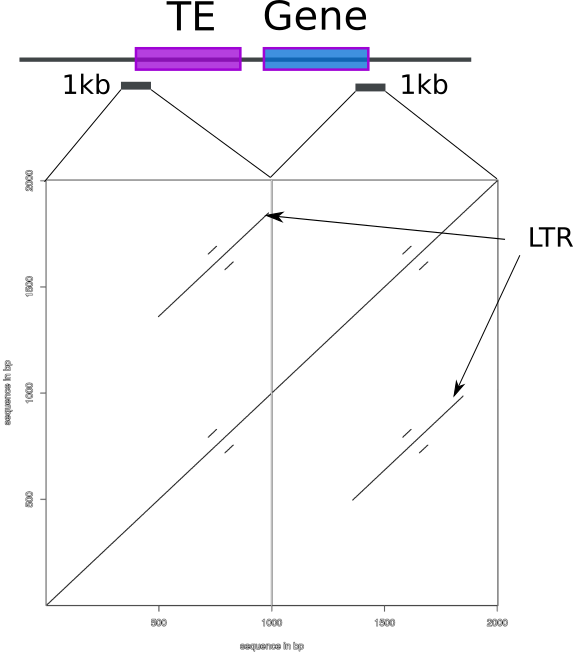

**Supplemental Figure 11:** Dot-plots of the end of insertion B. LTR repeats can be detected on each side of the insertion. Cf. **Supplemental Figure 12**.

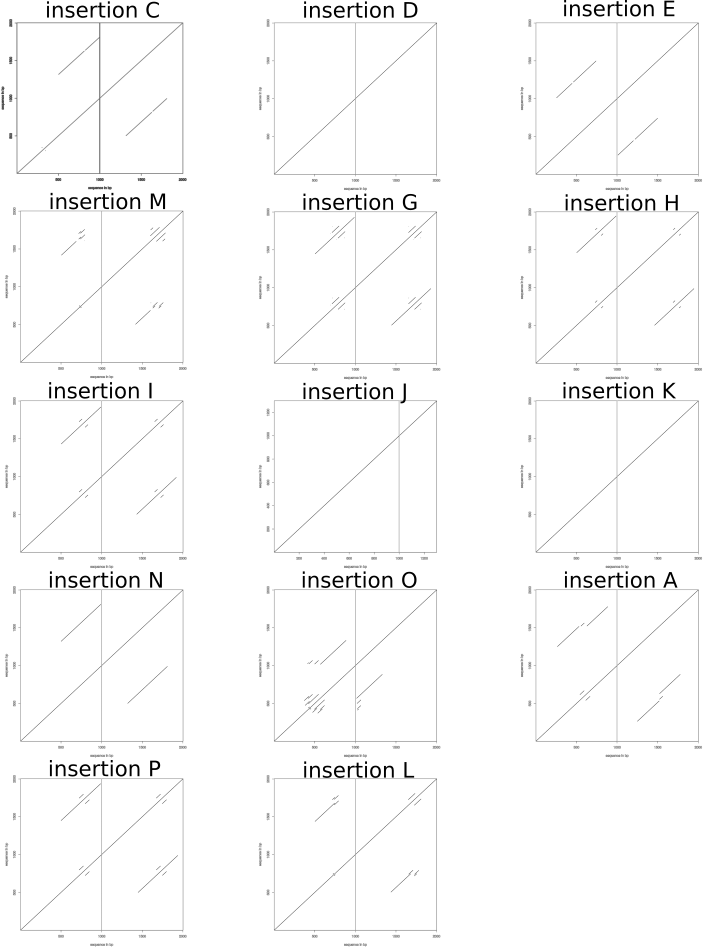

**Supplemental figure 12:** Dot plot of the ends of all insertions. Cf. **Supplemental Figure 11**.

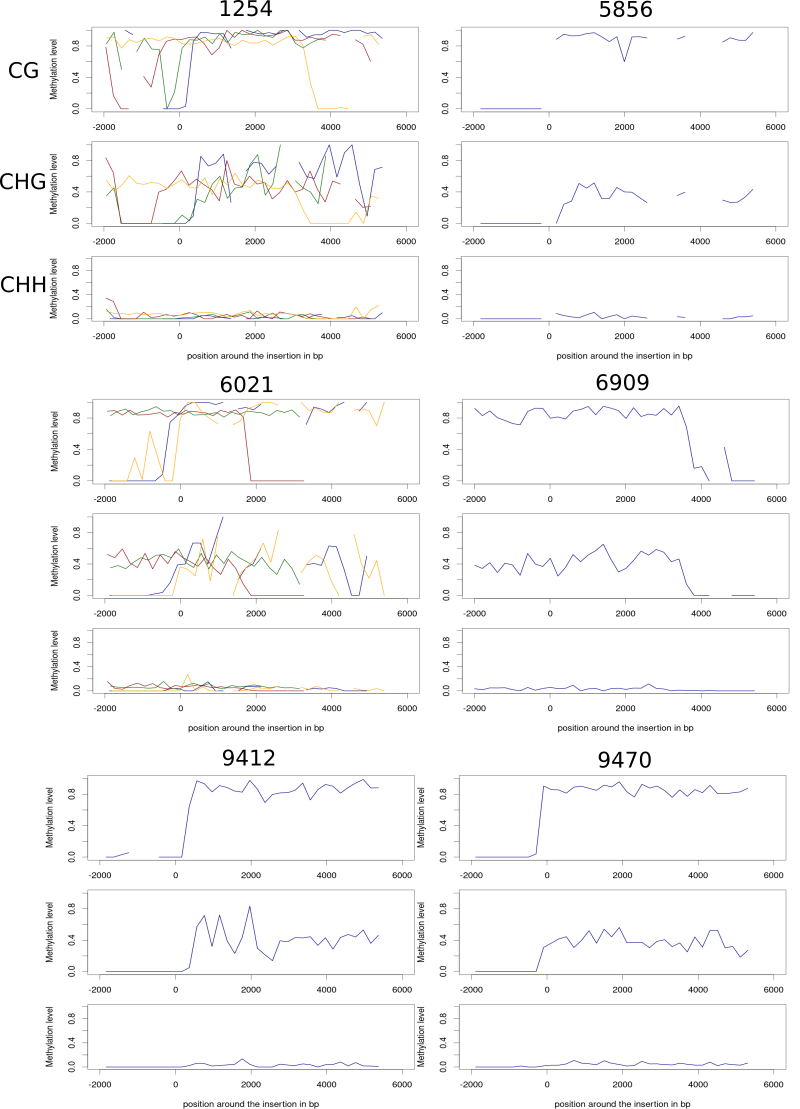

**Supplemental Figure 13:** Methylation profile across all copies of the TE+gene insertions. Bisulfite reads were mapped to the appropriate genomes and profiles extracted based on the inferred locations. Colors distinguish the different insertions found in each genome.

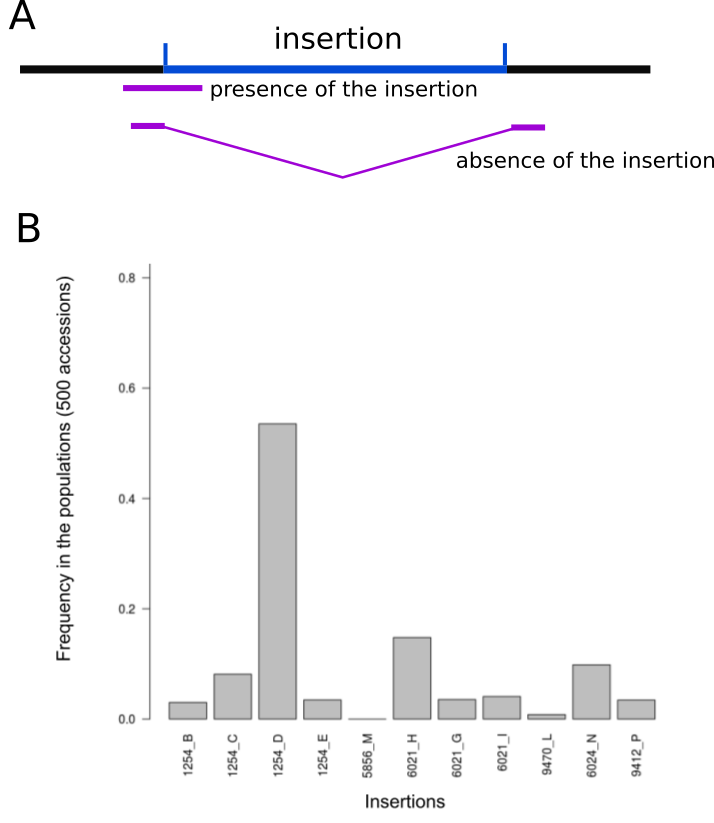

**Supplemental Figure 14:** Frequency of insertions of the new element in the 1001 Genomes. **(A)** Cartoon illustrating how the presence or absence of each known insertion in the seven full genomes was inferred using paired-end reads. **(B)** The population frequency of each insertion detected in the seven genomes.

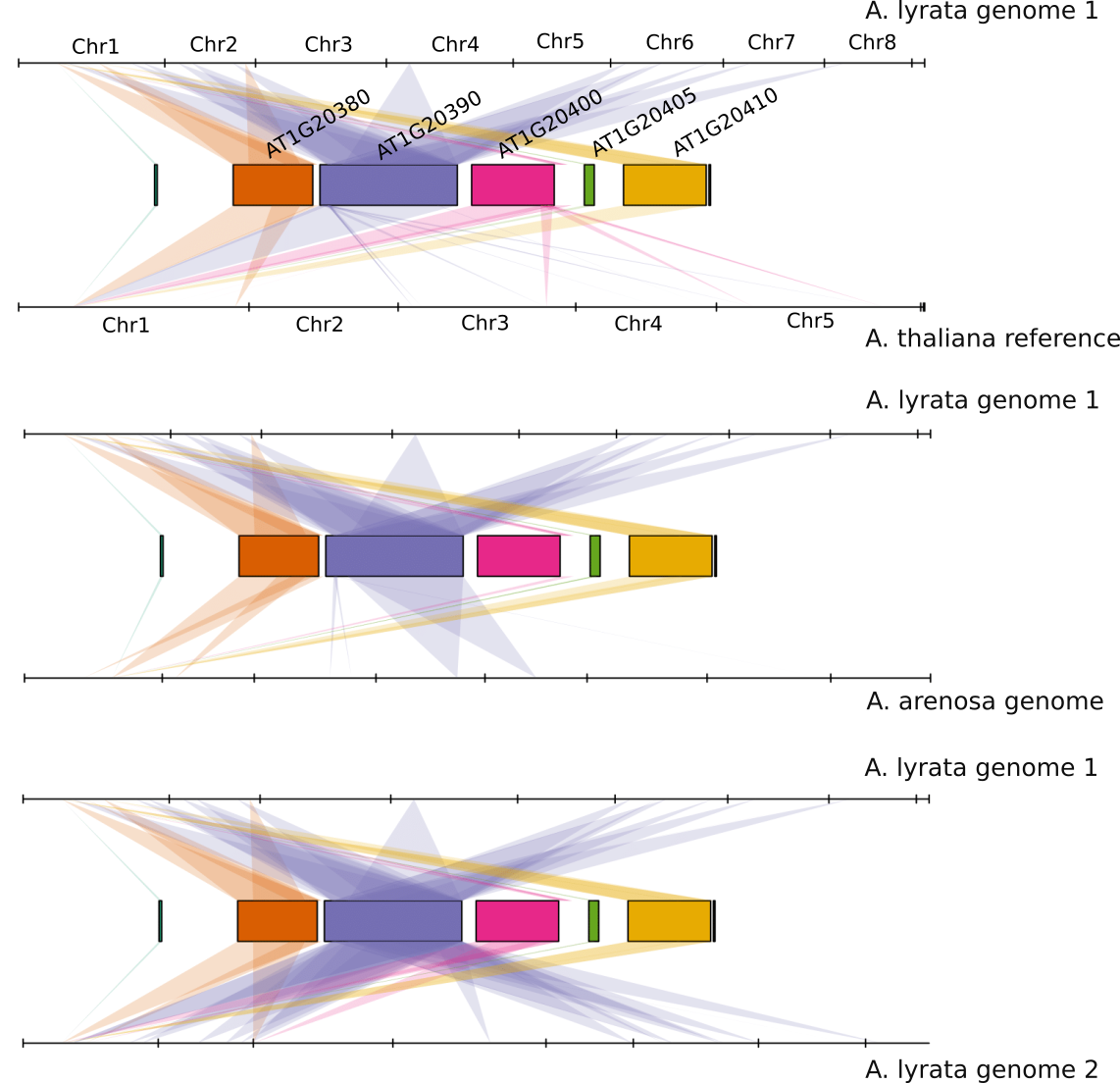

**Supplemental Figure 15:** Mapping of the AT1G20400 region in multiple species. The rectangle corresponds to annotated genes around AT1G20400 in the *A. thaliana* reference genome.

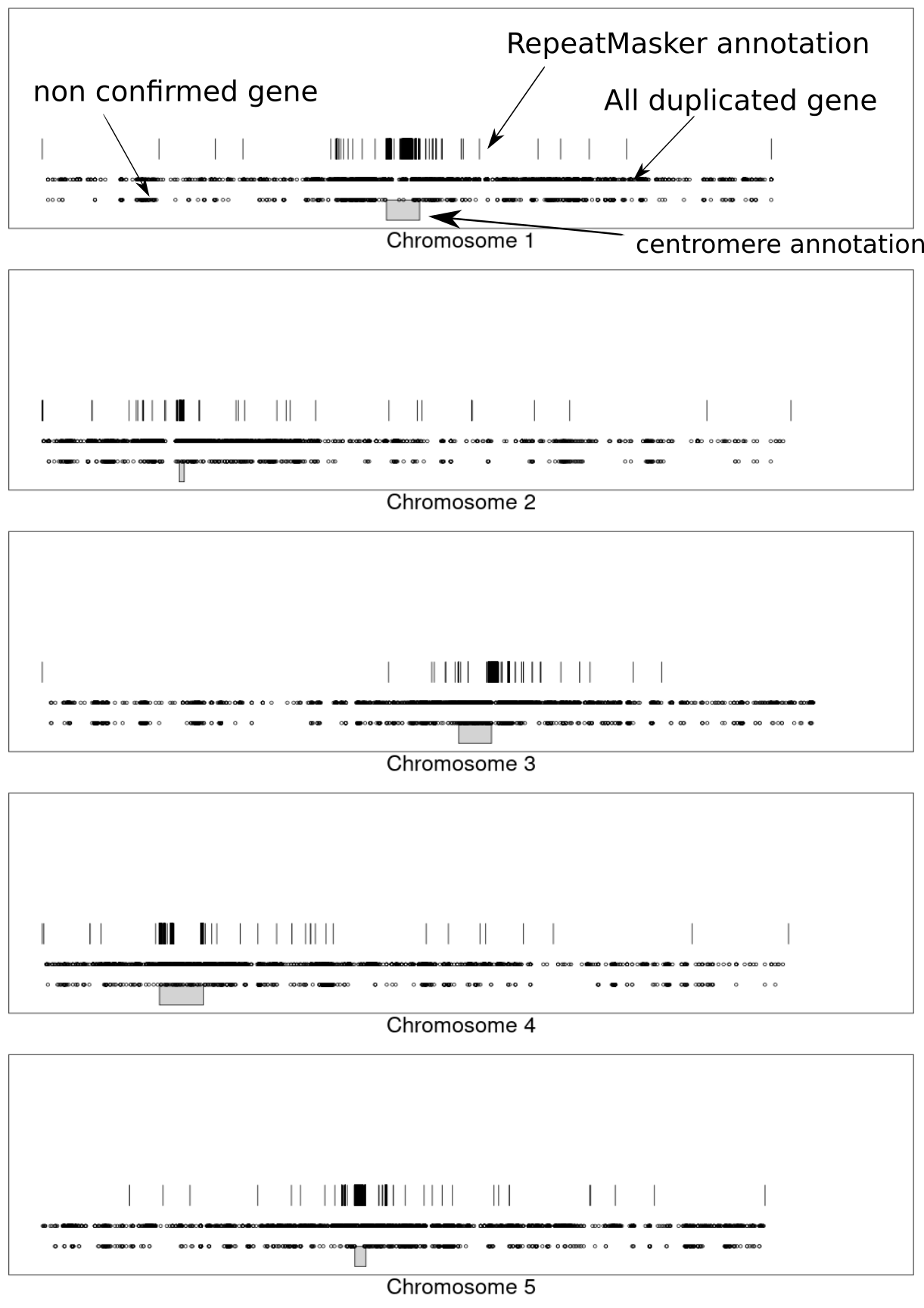

**Supplemental Figure 16:** Chromosomal position of putatively duplicated genes that could not be confirmed using full genome sequence. The figure shows the distribution of these genes compared to the distribution of all genes, and RepeatMasker annotation.

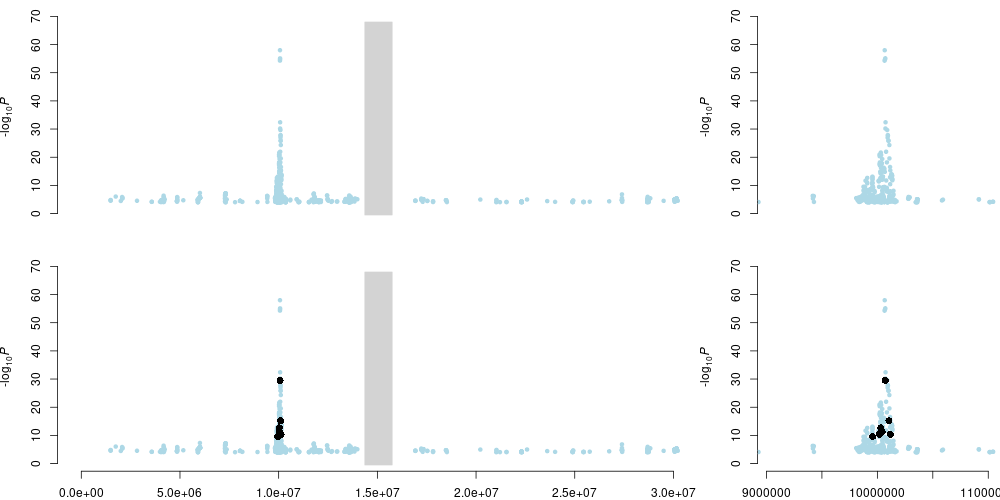

**Supplemental Figure 17**: Illustration of GWAS peak calling. The p-values from GWAS were used to run the filterFFT function from the NucleR package in R, using 0.05 for pcKeepComp option. To detect the peak, the function peakDetection was used with threshold=90%, width=1. The upper plot represents the -log10 p value of a representative GWAS of each SNPs for chromosome 1, and the lower plot represents the detected peak (black dots).

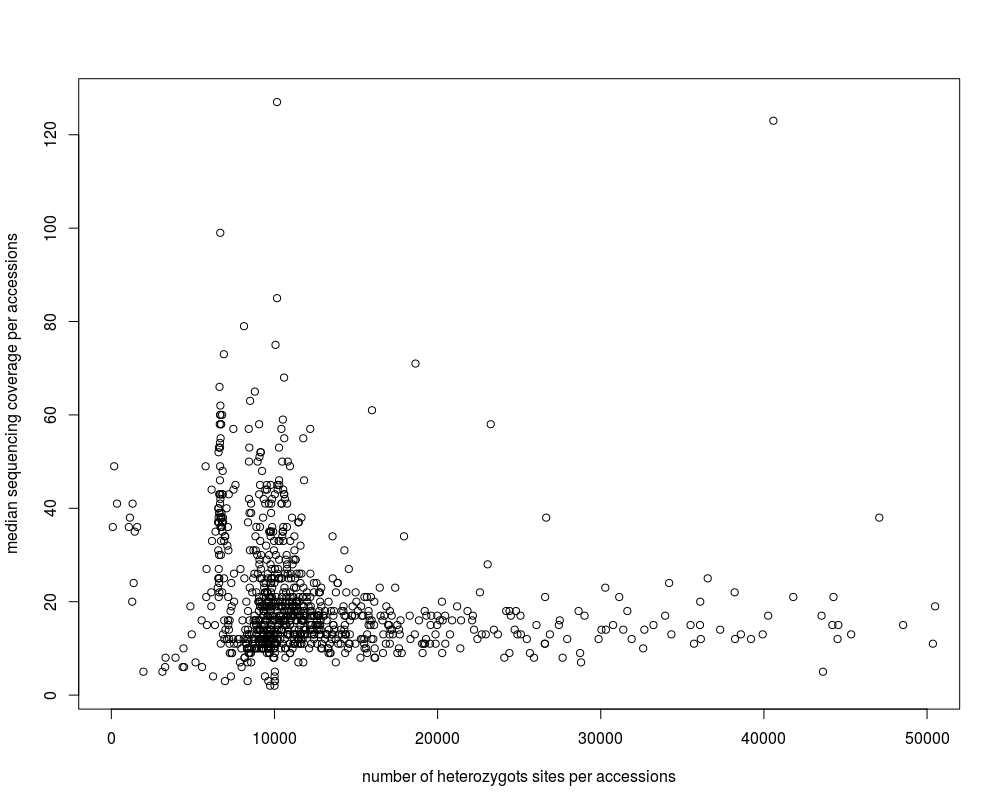
**Supplemental Figure 18:** The relationship between the number of pseudo-SNPs and sequencing coverage.. Each dot represents an accession.
